# Identification of a novel AP2 transcription factor in zygotes with an essential role in *Plasmodium* ookinete development

**DOI:** 10.1101/2022.04.11.487829

**Authors:** Tsubasa Nishi, Izumi Kaneko, Shiroh Iwanaga, Masao Yuda

## Abstract

The sexual phase of *Plasmodium* represents a crucial step in malaria transmission, during which these parasites fertilize and form ookinetes to infect mosquitoes. *Plasmodium* development after fertilization is thought to proceed with female-stored mRNAs until the formation of a retort-form ookinete; thus, transcriptional activity in zygotes has previously been considered quiescent. In this study, we reveal the essential role of transcriptional activity in zygotes by investigating the function of a newly identified AP2 transcription factor, AP2-Z. *ap2-z* was previously reported as a female transcriptional regulator gene whose disruption resulted in developmental arrest at the retort stage of ookinetes. In this study, although *ap2-z* was transcribed in females, we show that it was translationally repressed by the DOZI complex and translated after fertilization with peak expression at the zygote stage. ChIP-seq analysis of AP2-Z shows that it binds on specific DNA motifs, targeting the majority of genes known as an essential component of ookinetes, which largely overlap with the AP2-O targets, as well as genes that are unique among the targets of other sexual transcription factors. The results of this study also indicate the existence of a cascade of transcription factors, beginning with AP2-G, that proceeds from gametocytogenesis to ookinete formation.

**Author summary:** Sexual development in *Plasmodium* parasites, a causative agent of malaria, is essential for their transmission from vertebrate hosts to mosquitoes. This important developmental process proceeds as follows: formation of a gametocyte/gamete, fertilization and conversion of the zygote into the mosquito midgut invasive stage, called the ookinete. As a target of transmission blocking strategies, it is important to understand the mechanisms regulating *Plasmodium* sexual development. In this study, we assessed transcriptional regulation after fertilization by investigating the function of a novel transcription factor, AP2-Z. The results revealed the essential role of *de novo* transcription activated by AP2-Z in zygotes for promoting ookinete development. As transcriptional activity during the zygote stage has previously been considered silent in *Plasmodium*, novel genes important for ookinete formation can now be explored in the target genes of AP2-Z. Investigating the functions of these genes can help us understand the mechanisms of *Plasmodium* zygote/ookinete development and identify new targets for transmission blocking vaccines.

## Introduction

Malaria is a serious infectious disease caused by *Plasmodium* parasites of the Apicomplexa phylum, which are propagated among humans through bites from Anopheles mosquitoes [1]. Successful transmission from vertebrate hosts to mosquitoes requires these parasites to proceed with complex sexual development [2, 3]; therefore, understanding the mechanism of *Plasmodium* sexual development is a crucial aspect of malaria epidemiology.

*Plasmodium* sexual development begins when a subpopulation of asexual blood-stage parasites differentiate into male and female gametocytes in the host blood stream [4, 5]. Taken up through mosquito blood feeding, these gametocytes then develop into gametes and fertilize. The fertilized female gametes, or zygotes, then develop into a banana-shaped motile stage known as ookinetes, which then invade the midgut epithelia to proceed with further development beneath the basal lamina [6].

*Plasmodium* sexual development is regulated at both transcriptional and translational levels. At the beginning of sexual development, gametocytogenesis is induced by the AP2 transcription factor AP2-G, which is expressed in a subpopulation of blood-stage parasites [7–10]. Subsequently, the transcriptional repressor AP2-G2 starts to be expressed in the early stage and supports differentiation to gametocytes [11, 12]. Early gametocytes then differentiate into male and female gametocytes. For females, two transcription factors, AP2-FG and AP2-O3, function as a transcriptional activator and repressor, respectively, to promote development of female gametocytes [13, 14]. In addition, during female development, parasites prepare for development after fertilization by preserving mRNAs [15]. This preservation of mRNAs is conducted by a translational repressor complex, in which at least two factors, ATP-dependent RNA helicase DDX6 (DOZI) and trailer hitch homolog (CITH), are currently known to be involved [16–19].

In *Plasmodium berghei*, fertilized females undergo meiosis within 4 h post-fertilization [18] then start forming an apical protrusion approximately 8 h post-fertilization to develop into ookinetes [20]. For these processes, female-stored mRNA plays an essential role; parasites that lack *dozi* are not able to begin either meiosis or apical protrusion formation [15, 18]. In contrast, the role of *de novo* transcripts in zygotes has not been investigated, although it has been shown that *Plasmodium* zygotes can develop into retort-form ookinetes even in the presence of a transcriptional inhibitor [19]. Therefore, during the development of *Plasmodium* zygotes, transcriptional activity is considered to be quiescent, similar to animal embryos whose transcriptional activity is quiescent during the early stage after fertilization [21, 22]. For the late stage of ookinete development, parasites must activate *de novo* transcription. AP2-O is one transcription factor that is expressed from retort-form ookinetes to mature ookinetes [23], which activates the majority of known ookinete genes; thus, disruption of *ap2-o* results in impaired development of ookinetes [24]. In addition, AP2-O2 is also essential for ookinete maturation in *P. berghei* [25], although disruption of *ap2-o2* in *P. yoelii* does not affect ookinete development [26]. Thus, according to existing studies of transcriptional and translational regulators, *Plasmodium* ookinete development is currently considered to be promoted by female-stored mRNAs in the early stage and *de novo* transcripts in the late stage.

In a previous study, we identified an AP2 transcription factor-related gene, *ap2r-1*, as a target gene of AP2-G and AP2-FG, and showed that its disruption affects ookinete development [10]. Here, we demonstrate that AP2R-1 is an AP2 transcription factor that functions in zygotes; hence, we rename it AP2-Z. By investigating the functions of AP2-Z, we reveal the essential role of *de novo* transcription in zygotes for ookinete formation. Moreover, we also demonstrate the existence of a transcriptional cascade, which starts with AP2-G, during *Plasmodium* development from gametocytes to ookinetes.

## Results

### *ap2r-1* is an AP2-family protein gene conserved in Apicomplexan parasites

We previously identified *ap2r-1* (PBANKA_0612400) as a target gene of AP2-G and AP2-FG. This gene is expressed in female gametocytes, and its disruption results in arrested zygote development at the retort stage of ookinetes. AP2R-1 has a putative AP2 domain that has not been identified by homology searches because of a non-canonical linker peptide between the second and third sheet (Fig 1A). The domain is highly conserved in the *Plasmodium* species except for the linker peptide, which varies in amino acid sequence and length among the species. To investigate whether AP2R-1 is conserved in other organisms, we searched for any proteins with a domain similar to the putative AP2 domain of AP2R-1. Protein–protein BLAST (blastp) search identified proteins with a homologous domain to the putative AP2 domain of AP2R-1 from Apicomplexan parasites, such as *Toxoplasma gondii* and *Eimeria tenella* (Fig 1B). The conserved amino acids among *Plasmodium* AP2R-1 and these blastp-searched proteins were mostly found within the putative beta sheets and alpha helix, and did not have the non-canonical linker peptide, except for the protein from *Hepatocystis* (Fig 1B). Notably, these conserved amino acids contained consensus amino acids for the AP2 domain [27], indicating that this conserved domain is actually an AP2 domain. The homologous protein from *Toxoplasma* is consistently annotated as “AP2 domain transcription factor AP2X-6,” and the homologous domain from all blastp-searched proteins without the non-canonical linker was assigned as “AP2” in the SMART analysis. The phylogenetic tree of *Plasmodium* AP2R-1 and the blastp-searched proteins was topologically consistent with the species tree of Apicomplexa, which further suggested that these proteins are orthologs (Fig 1C). In this regard, their ancestral protein probably did not include the non-canonical linker peptide within the conserved domain, with *Plasmodium* and *Hepatocystis* species acquiring this linker during evolution instead. Collectively, we concluded that AP2R-1 is an AP2-family protein conserved in the phylum Apicomplexa.

**Fig 1.**
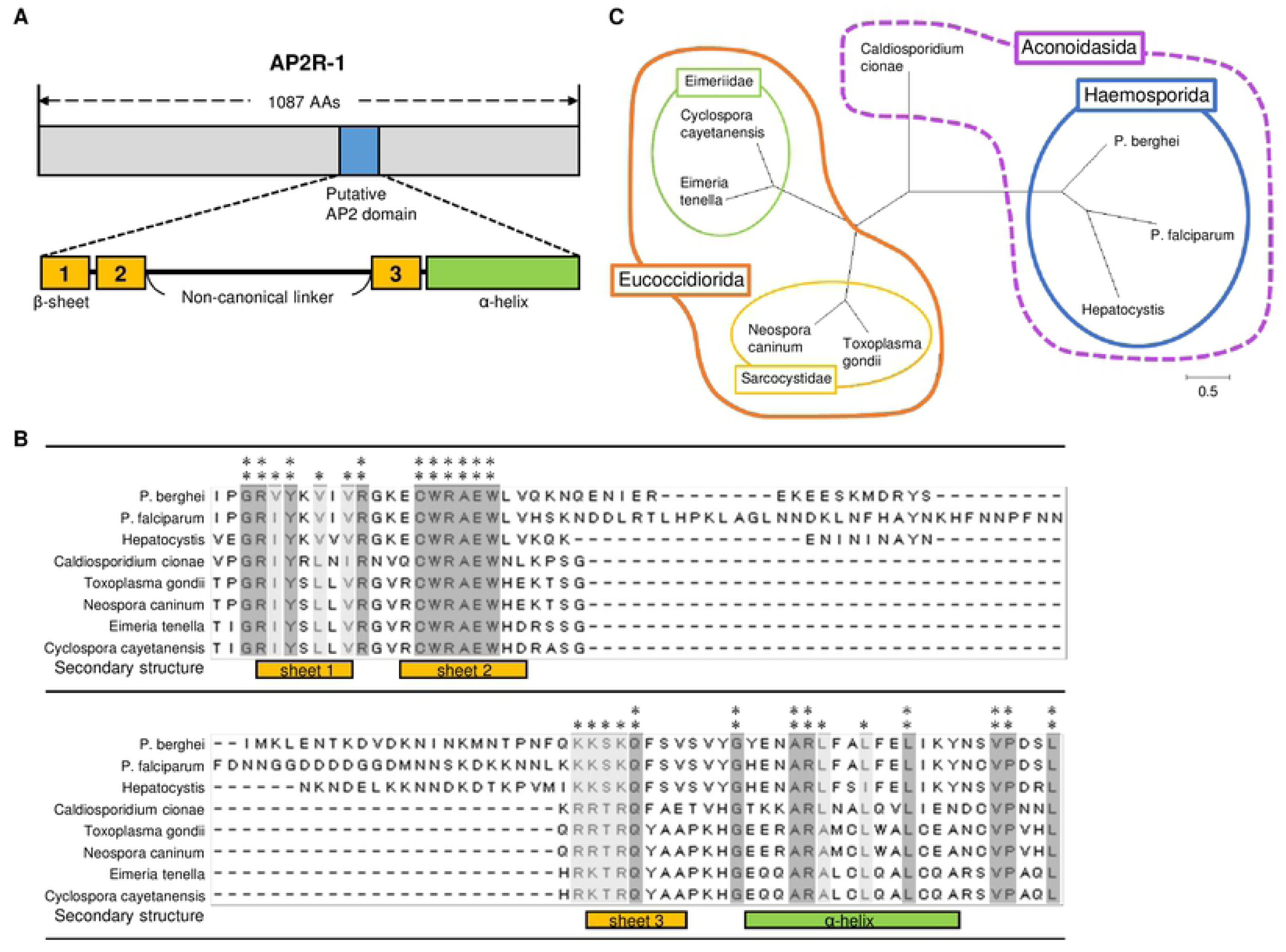
Identification of possible AP2R-1 orthologs in Apicomplexan parasites. (A) A schematic illustration of AP2R-1. Blue box shows the AP2 domain. Beta-sheet and α-helix regions in the AP2 domain were predicted using PSIPRED (http://bioinf.cs.ucl.ac.uk/psipred/) and indicated in orange and green boxes, respectively. (B) Alignment of conserved amino acid sequences from AP2R-1 and blastp-searched proteins by the ClustalW program in Mega X. Positions at which all sequences have an identical amino acid are indicated by two asterisks, whereas positions with amino acid residues of the same property are indicated by one asterisk. Amino acid sequences were retrieved from PlasmoDB or the NCBI database (*P*. *berghei*, PBANKA_0612400; *P*. *falciparum*, PF3D7_0411000; *Hepatocystis*, VWU49277; *Cardiosporidium cionae*, KAF8822898; *Toxoplasma gondii*, XP_018635762; *Neospora caninum*, XP_003884720; *Eimeria tenella*, XP_013229162; *Cyclospora cayetanensis*, OEH74626). (C) Phylogenetic tree of AP2R-1 and blastp-searched proteins, which was inferred from their whole amino acid sequences using the Maximum Likelihood method and JTT matrix-based model. Tree is drawn to scale, with branch lengths measured according to the number of substitutions per site. Species of the same family, order, and class are enclosed by thin, thick, and dashed lines, respectively.

### *ap2r-1* is translationally repressed in female gametocytes by the DOZI complex and expressed in zygotes

In our previous study, we tagged AP2R-1 with mNeonGreen fluorescent protein (mNG) using a conventional homologous recombination method (AP2R-1::mNG), and observed a fluorescent signal in the nucleus of female gametocytes, consistent with the previous observation that it is a target of AP2-G and AP2-FG [10]. To evaluate whether *ap2r-1* is actually transcribed by AP2-G and AP2-FG, we planned to perform a promoter assay on the endogenous *ap2r-1*. However, when we developed parasites expressing green fluorescent protein (GFP)-fused AP2R-1 (AP2R-1::GFP) by the CRISPR/Cas9 system using the Cas9-expressing parasite (PbCas9) [28] (Fig S1A) to perform the promoter assay, we observed no fluorescent signal in any blood-stage parasites, including female gametocytes (Fig 2A).

**Fig 2.**
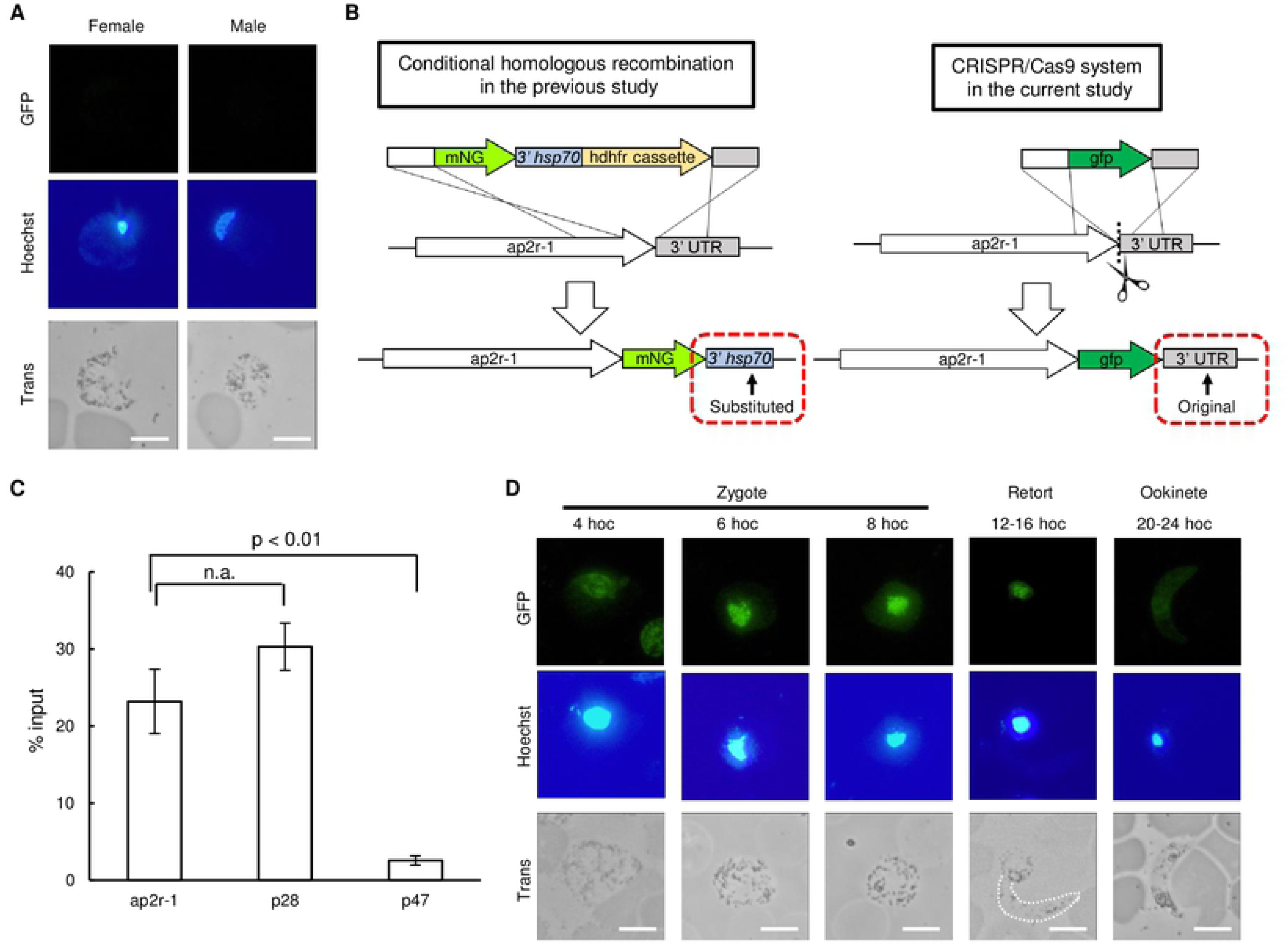
Expression profile of AP2R-1 during sexual development and its translational repression in females. (A) Expression of AP2R-1 during the blood stage using the AP2R-1::GFP parasite. Nuclei were stained with Hoechst 33342. Scale bar = 5 μm. (B) Schematic illustrations of fluorescent protein-tagging strategies of AP2R-1. In the AP2R-1::mNG parasite, the 3’ UTR of *ap2r-1* was substituted upon fusion with *mNG* (left). In the AP2R-1::GFP parasite, the endogenous 3’UTR was retained upon *gfp*-fusion performed by CRISPR/Cas9 (right). (C) Enrichment of DOZI-associated transcripts evaluated by RNA immunoprecipitation- qPCR analysis. Pre-immunoprecipitated samples were used as the input. Enrichment of *p28* and *p47* transcripts was analyzed as a positive and negative control, respectively. The *p*-value on the graph was calculated by Student’s t-test. Error bars indicate the standard error of the mean (n = 3). (D) Expression of AP2R-1 during ookinete culture using the AP2R-1::GFP parasite. Nuclei were stained with Hoechst 33342. Apical protrusion of the retort-form ookinete at 16 hoc is highlighted with a white dashed line. Scale bar = 5 μm.

In the homologous recombination method used previously, the 3’ untranslated region (UTR) of *ap2r-1* was replaced by that of *hsp70* upon tagging AP2R-1 with mNG (Fig 2B, left) [10]. However, in this study, we inserted *gfp* at the 3’ end of *ap2r-1* by CRISPR/Cas9 without replacing the original 3’ UTR (Fig 2B, right). In female gametocytes of the *Plasmodium* species, a subset of genes is translationally repressed by the DOZI complex in a 5’ UTR- or 3’ UTR-dependent manner [17, 19]. Therefore, we assumed that *ap2r-1* is translationally repressed by the DOZI complex in normal females but translated in females of AP2R-1::mNG because of the replacement of 3’ UTR. Consistent with this assumption, strong fluorescence was observed in the nucleus of AP2R-1::GFP zygotes.

To further address whether *ap2r-1* is translationally repressed by the DOZI complex, we developed parasites expressing GFP-fused DOZI (DOZI::GFP) and performed RNA immunoprecipitation (RIP) followed by reverse transcription quantitative polymerase chain reaction (RT-qPCR) analysis. The analysis showed that more than 20% of *ap2r-1* transcript was immunoprecipitated with anti-GFP antibody (Fig 2C). The percentage was comparable to that of *p28*, which is known as a target of the DOZI complex [15, 19]. On the other hand, the transcript of *p47*, which is a gene translated in females [29], was only slightly detected in the immunoprecipitated samples. Therefore, these results validated that *ap2r-1* was translationally repressed by the DOZI complex in female gametocytes.

Given the above results, we next assessed the expression of AP2R-1 in a time course manner during zygote/ookinete development. Nuclear-localized fluorescence began to appear in zygotes at 4 h after the start of ookinete culture (hoc) and became stronger at 6 hoc (Fig 2D). The strong fluorescence continued until zygotes started forming an apical protrusion (8–12 hoc). The signal then faded in retort-form ookinetes and became barely detectable in the nuclei of banana-shaped ookinetes (Fig 2D). Therefore, AP2-Z is mainly expressed in zygotes prior to apical protrusion formation. Considering the expression of AP2-O in the late stage of ookinete development [23], these results might indicate that AP2-Z is responsible for transcriptional regulation until AP2-O starts to be expressed. As a result, we renamed *ap2r-1* as *ap2-z* (and renamed AP2R- 1::GFP as AP2-Z::GFP).

### *ap2-z* is transcribed by AP2-G and AP2-FG in female gametocytes

To evaluate whether *ap2-z* transcription is activated downstream of AP2-G and AP2-FG, we disrupted their binding motifs located upstream of *ap2-z* by CRISPR/Cas9 using AP2-Z::GFP parasites (AP2-Z::GFP_cismut) (Fig 3A and S1B) and assessed the expression of AP2-Z in the zygotes. The upstream region of *ap2-z* has two binding motifs of AP2-G [10] and one motif of AP2-FG [13]. These motifs are located near to each other, with one of the AP2-G motifs overlapping with the AP2-FG motif, at approximately 650 bp upstream from the start codon of *ap2-z* (Fig 3A). Disruption of these three motifs resulted in complete loss of fluorescence in the nucleus of zygotes (Fig 3B). Furthermore, the AP2-Z::GFP_cismut parasites lost the ability to produce banana-shaped ookinetes, as with *ap2-z* knockout parasites [*ap2-z*(-)] (Fig 3C and 3D). We further examined the expression of *ap2-z* in the AP2-Z::GFP_cismut at the RNA level. Mice infected with AP2-Z::GFP or AP2-Z::GFP_cismut parasites were treated with sulfadiazine to enrich gametocytes by killing asexual blood-stage parasites. Total RNA was then harvested from the infected blood and subjected to RT-qPCR analysis. The amount of *ap2-z* transcripts relative to that of *p28* was downregulated more than 200-fold in AP2-Z::GFP_cismut compared to that in AP2-Z::GFP (Fig 3E). Collectively, these results demonstrated that transcription of *ap2-z* is activated by AP2-G and AP2-FG in the early and female gametocyte, respectively.

**Fig 3.**
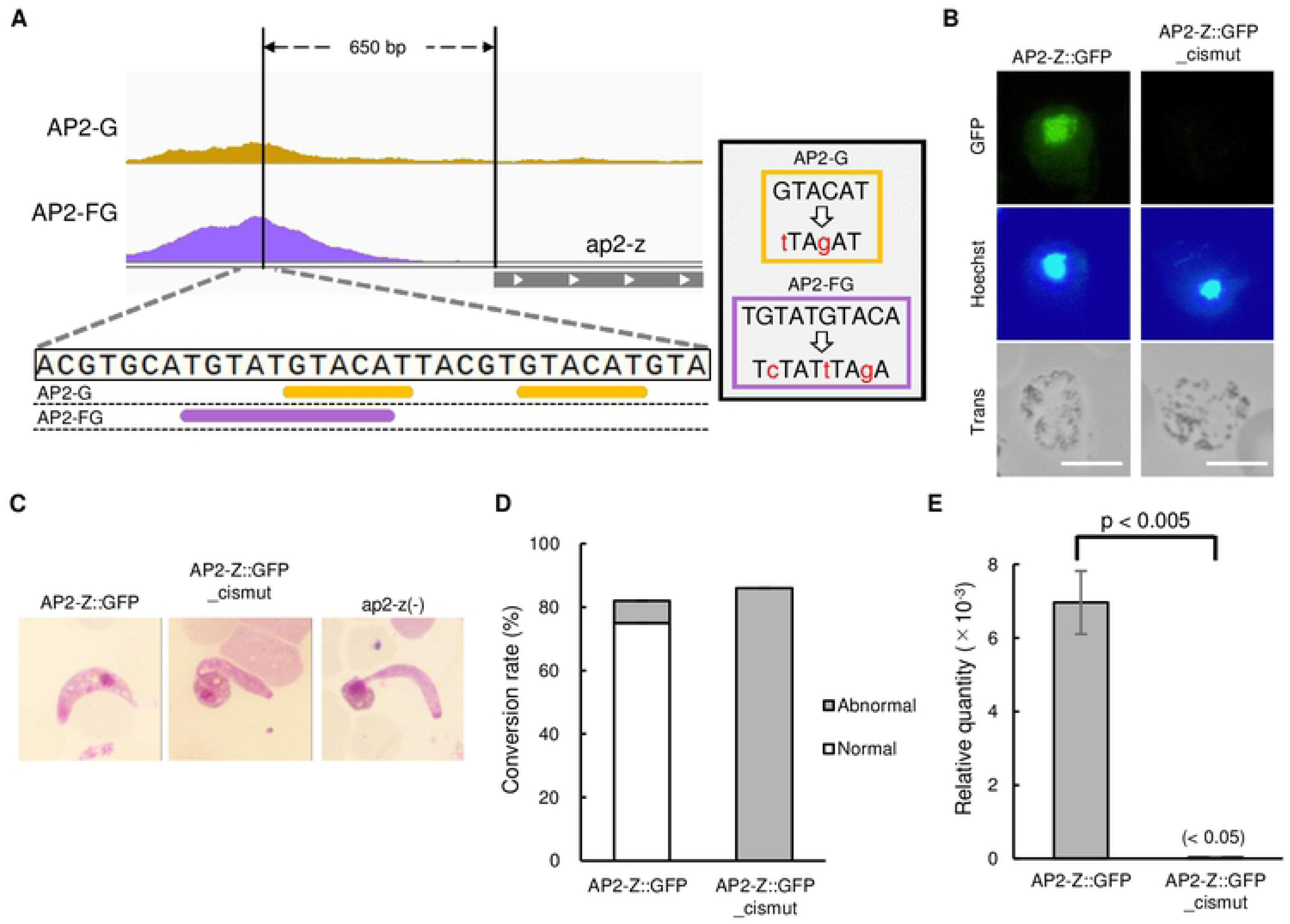
Transcriptional activation of *ap2-z* by AP2-G and AP2-FG. (A) Integrative Genomics Viewer images from the ChIP-seq data of AP2-G and AP2-FG for the upstream region of *ap2-z*. Binding motifs of AP2-G and AP2-FG in the peak region are indicated by orange and purple bars, respectively. Mutations introduced to these motifs are described on the right. (B) Expression of AP2-Z in the AP2-Z::GFP_cismut at 6 hoc. Nuclei were stained with Hoechst 33342. Scale bar = 5 μm. (C) Representative Giemsa-stained image of ookinete in AP2-Z::GFP, AP2-Z::GFP_cismut, and *ap2-z*(-) parasites at 20 hoc. (D) Rate of conversion to ookinetes against all female-derived cells in AP2-Z::GFP and AP2-Z::GFP_cismut at 20 hoc. (E) Relative amounts of *ap2-z* transcripts in gametocyte assessed by RT-qPCR analysis. The *p28* gene was used as the internal control. Value for AP2-Z::GFP_cismut is given in brackets as it is too small to be depicted in the graph. The *p*-value on the graph was calculated by the Student’s t-test. Error bars indicate the standard error of the mean (n = 3).

### AP2-Z binds to specific sites of the genome recognizing (T/C)(A/C)TG(A/T)AC(A/G) motifs

To assess the target genes of AP2-Z, we performed chromatin immunoprecipitation followed by high-throughput sequencing (ChIP-seq) analysis using AP2-Z::GFP parasites. The parasites were harvested at 6 hoc and subjected to ChIP-seq experiments using anti-GFP antibody, and the experiments were performed in duplicate. The overall peak patterns were similar between each replicate, indicating good reproducibility of the two experiments (Fig 4A). Experiment 1 and 2 identified 1,214 and 1,256 peaks with fold enrichment > 4.0, respectively, with 1,110 overlapping peaks between the two experiments (Fig 4B, Table S1A and S1B).

**Fig 4.**
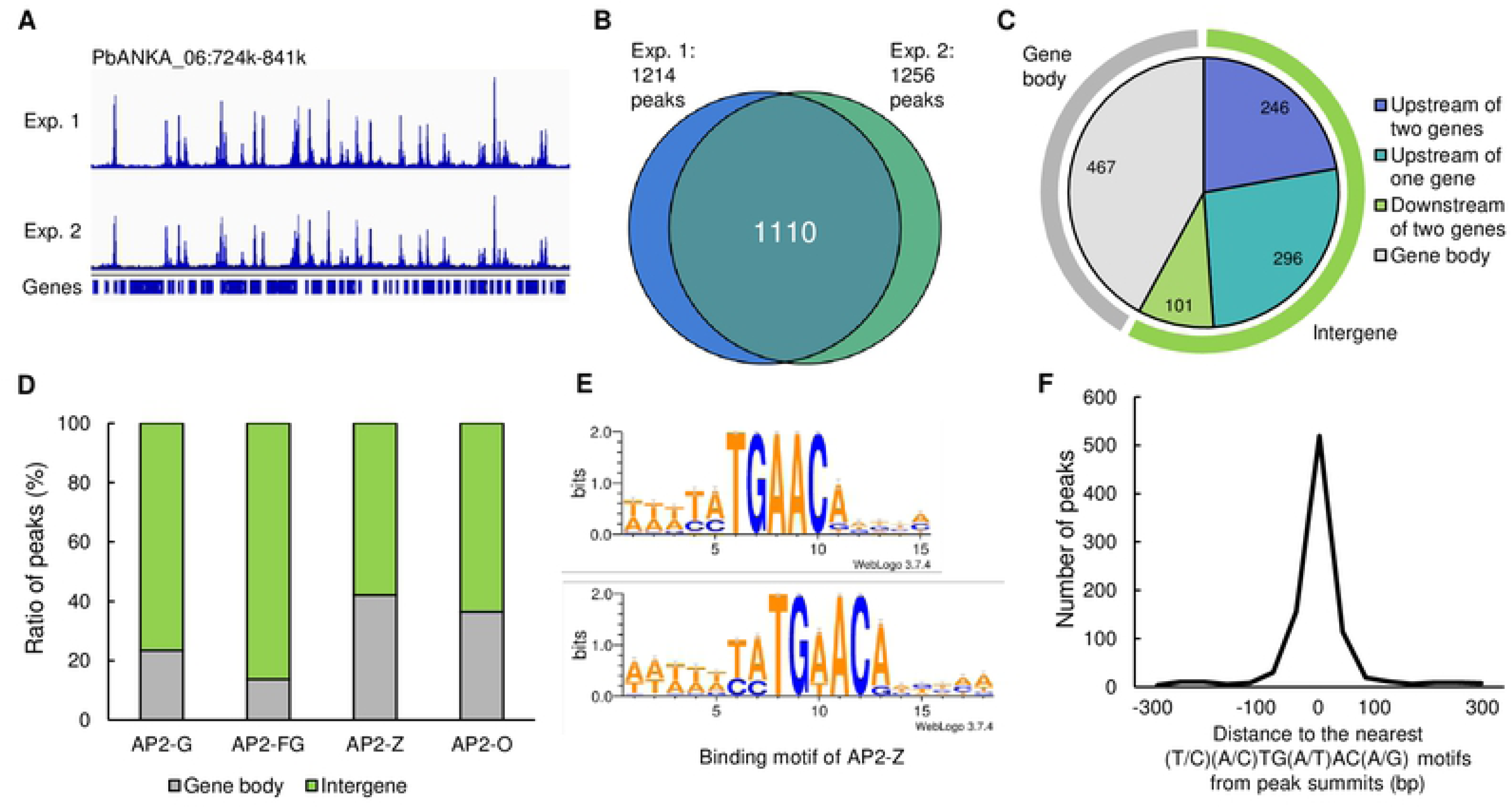
ChIP-seq analysis of AP2-Z. (A) Integrative Genomics Viewer images from ChIP-seq experiment 1 and 2 of AP2-Z in a part of chromosome 6. Histograms show sequence coverage of ChIP data at each base. (B) Venn diagram of number of peaks identified in ChIP-seq experiment 1 and 2. (C) Classification of peak locations. Light green and gray bars around the pie chart indicate the intergenic region and gene body region, respectively. (D) Ratio of peaks located on the gene body and intergenic region for ChIP-seq of AP2-G, AP2-FG, AP2-Z, and AP2-O. (E) Sequence logo constructed from TGAAC motifs in the peak regions (top) and predicted binding motif of AP2-Z (bottom). Sequence logos are depicted using WebLogo3 (http://weblogo.threeplusone.com/create.cgi). (F) Histogram showing the distance between the peak summit and the nearest (T/C)(A/C)TG(A/T)AC(A/G) motif for each ChIP peak.

Of the 1,110 common peaks, 643 peaks (57.9%) were located within intergenic regions, most of which were located upstream of the genes (Fig 4C). The ratio of peaks located on gene bodies (42.1%) was clearly higher than that in the ChIP-seq of AP2-G and AP2-FG, which were expressed in the gametocyte stages (Fig 4D) [10, 13]. However, when we analyzed our previous ChIP-seq data of AP2-O, which is an essential transcription factor for ookinete development, the ratio of peaks located on gene bodies was 36.5%, which was comparable to that of AP2-Z (Fig 4D) [24]. As both AP2-Z and AP2-O are expressed after parasites experience meiosis, these results might suggest that chromosomes are not fully assembled after meiotic replication during zygote/ookinete development; thus, gene bodies become relatively depleted for nucleosomes

Next, we searched for motifs enriched in the peak regions to predict the binding motif of AP2-Z. Motif enrichment analysis by Fisher’s exact test showed that the TATGAACA motif was most enriched within 50 bp from the peak summits, with a *p*- value of 3.69 × 10^-215^. We then identified enrichment of several further motifs that contained TGAAC and, taking them together, obtained the (T/C)(A/C)TGAAC(A/G) motif. When constructing the sequence logo by WebLogo3 using the sequences around the TGAAC motifs nearest to each peak summit, the above 8-bp motif was consistently depicted (Fig 4E, top). In addition to the (T/C)(A/C)TGAAC(A/G) motif, we also found enrichment of motifs that match (T/C)(A/C)TGTACA with a *p*-value of 5.16 × 10^-84^. Taken together, we obtained the (T/C)(A/C)TG(A/T)AC(A/G) motif as a putative binding motif of AP2-Z (Fig 4E, bottom). The motif was identified within 100 bp from the summits of 75.5% peaks; the closer the motif was to the summits, the more peaks were identified (Fig 4F).

### AP2-Z broadly targets genes that are important for ookinete development

Next, we identified the target genes of AP2-Z from the ChIP-seq data. The analysis showed that AP2-Z was bound within the 1,200-bp region upstream of 516 genes (Table S2). To evaluate the role of AP2-Z in zygote development, we classified these target genes into several groups according to their product descriptions annotated on PlasmoDB (https://plasmodb.org/plasmo/app/). The target genes contained 330 functionally annotated genes. Of these annotated target genes, 52 genes were categorized to the group ‘transcription and translation,’ which was the largest of 14 functional groups (Fig 5A). This group contained putative translation initiation factor genes, such as *eIF3b* and *eIF6*, elongation factor genes, and genes related to ribosomal biogenesis. Furthermore, the group ‘transcription and translation’ also included *ap2-o*, which is the gene coding a transcription factor essential for ookinete maturation (Fig 5B). In addition, although not identified as a target because of the 1,200-bp cutoff, we found a peak at approximately 1,500 bp upstream of *ap2-o2*. Considering the fact that *ap2-z* is activated by AP2-G and AP2-FG, these results suggested the existence of a cascade of transcription factors, starting from AP2-G, during the sexual development of *Plasmodium*. Besides *ap2-o* and *ap2-o2*, *ap2-z* itself was also included in the targets, suggesting the existence of transcriptional autoregulation (Fig 5B), a characteristic widely observed during prokaryotic and eukaryotic transcriptional regulation [30, 31]. As AP2-G and AP2-FG are no longer expressed after fertilization, this autoregulation is probably required for the stable expression of AP2-Z during zygote development.

**Fig 5.**
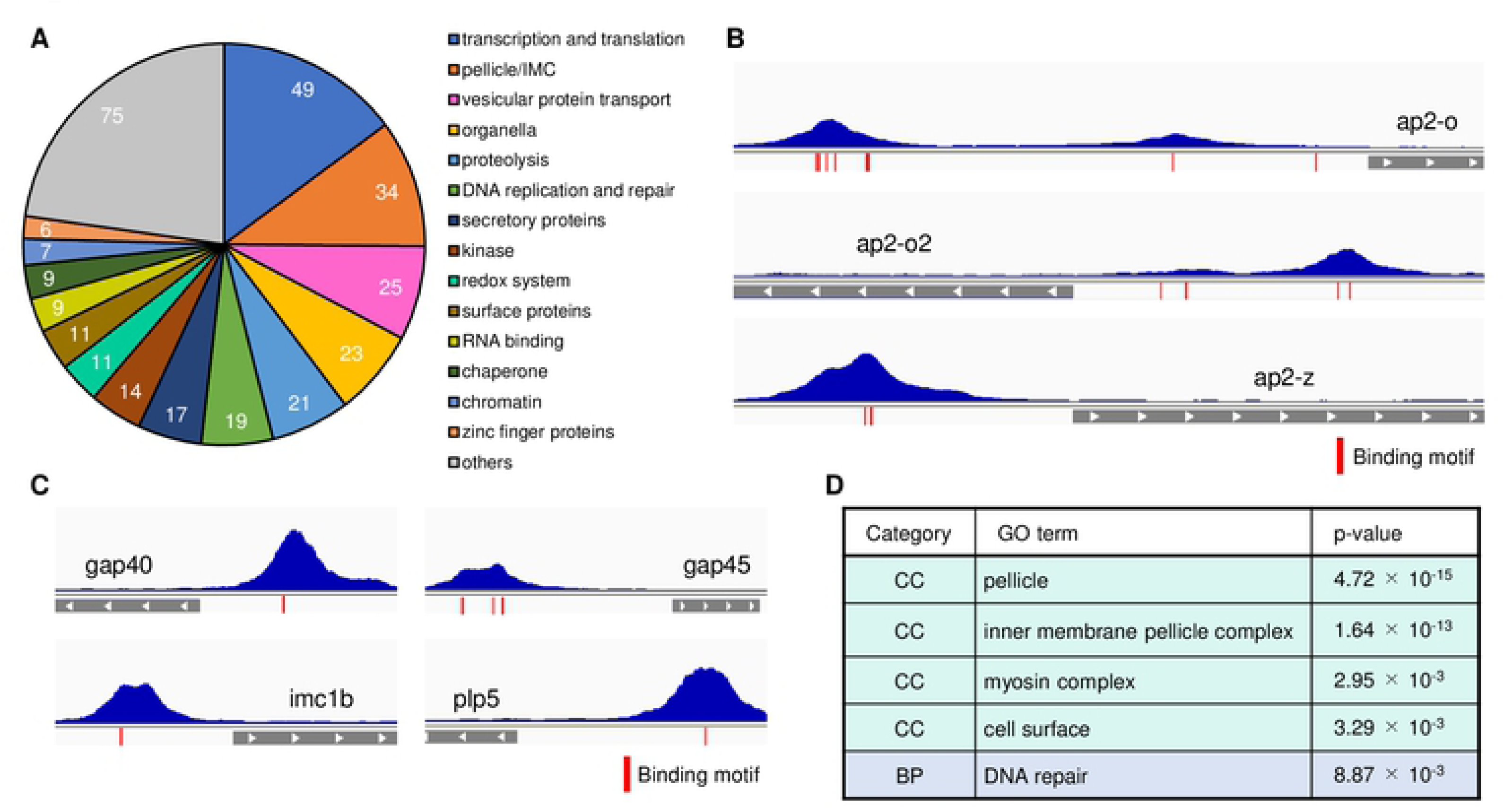
Target analysis of AP2-Z using ChIP-seq data. (A) Classification of the target genes of AP2-Z into 14 functional groups according to their product description annotated on PlasmoDB. (B) Transcription factor genes in the targets of AP2-Z. Histograms show sequence coverage of the ChIP sample in experiment 1. Gray bars indicate a partial open reading frame. Positions of the AP2-Z binding motifs are indicated in red. (C) Representative peak images of target genes essential for ookinete development. (D) GO analysis showing enrichment of genes with specific functions in the targets of AP2-Z. CC = Cellular Components, BP = Biological Process.

Other than the ‘transcription and translation’ group, the target genes largely contained genes that are important for ookinete development (Fig 5C). The group ‘pellicle/IMC’ included *gap40*, *gap45*, and 15 inner membrane complex (IMC) protein genes, which are related to the pellicle structure of ookinetes. Most of the known ookinete microneme genes, such as *plp3*, *celtos*, and *ctrp*, were included in the group ‘secretory proteins.’ Gene ontology (GO) analysis further supported these results. In the category ‘Cellular Components (CC)’, the target genes of AP2-Z were most enriched in ‘pellicle,’ with a *p*-value of 4.72 × 10^-15^ (Fig 5D). Furthermore, in the category ‘Biological Process (BP)’, the target genes were most enriched in ‘DNA repair,’ which indicated that AP2-Z may also contribute to genome stability after meiotic replication in the zygote stage.

### Target genes of AP2-Z are activated in early zygotes

Since *de novo* transcription in zygotes were previously considered silent, we next investigated whether the target genes of AP2-Z are actually transcribed in the zygotes. We performed RNA-seq analysis on the wild-type *Plasmodium berghei* ANKA strain (WT) at the gametocyte stage and 6 hoc to identify genes upregulated after fertilization. Compared to gametocytes, 512 genes were significantly upregulated (log_2_(fold change) > 1, *p*-value adjusted for multiple testing according to the Benjamini–Hochberg procedure (*p*-value adj) < 0.01) in the parasites at 6 hoc (Fig 6A, Table S3). These zygote- upregulated genes (ZUG) contained known ookinete genes, such as *cdpk3*, *celtos*, and *ctrp*, indicating that transcription of these ookinete genes already occurs during the early stage of zygote development. Of the 512 ZUGs, 198 genes were targets of AP2-Z, showing significant enrichment with a *p*-value of 3.7 × 10^-69^ by Fisher’s exact test. Furthermore, more than 80% of the target genes were more highly expressed in zygotes at 6 hoc than in gametocytes (log_2_(fold change) > 0) (Fig 6B), indicating that the targets of AP2-Z were transcribed after fertilization. Detecting significant upregulation in only a subset of the target genes (38%) was probably due to the overlap with female-stored mRNAs. Collectively, these results indicated that the target genes of AP2-Z are actively transcribed during the zygote stage.

**Fig 6.**
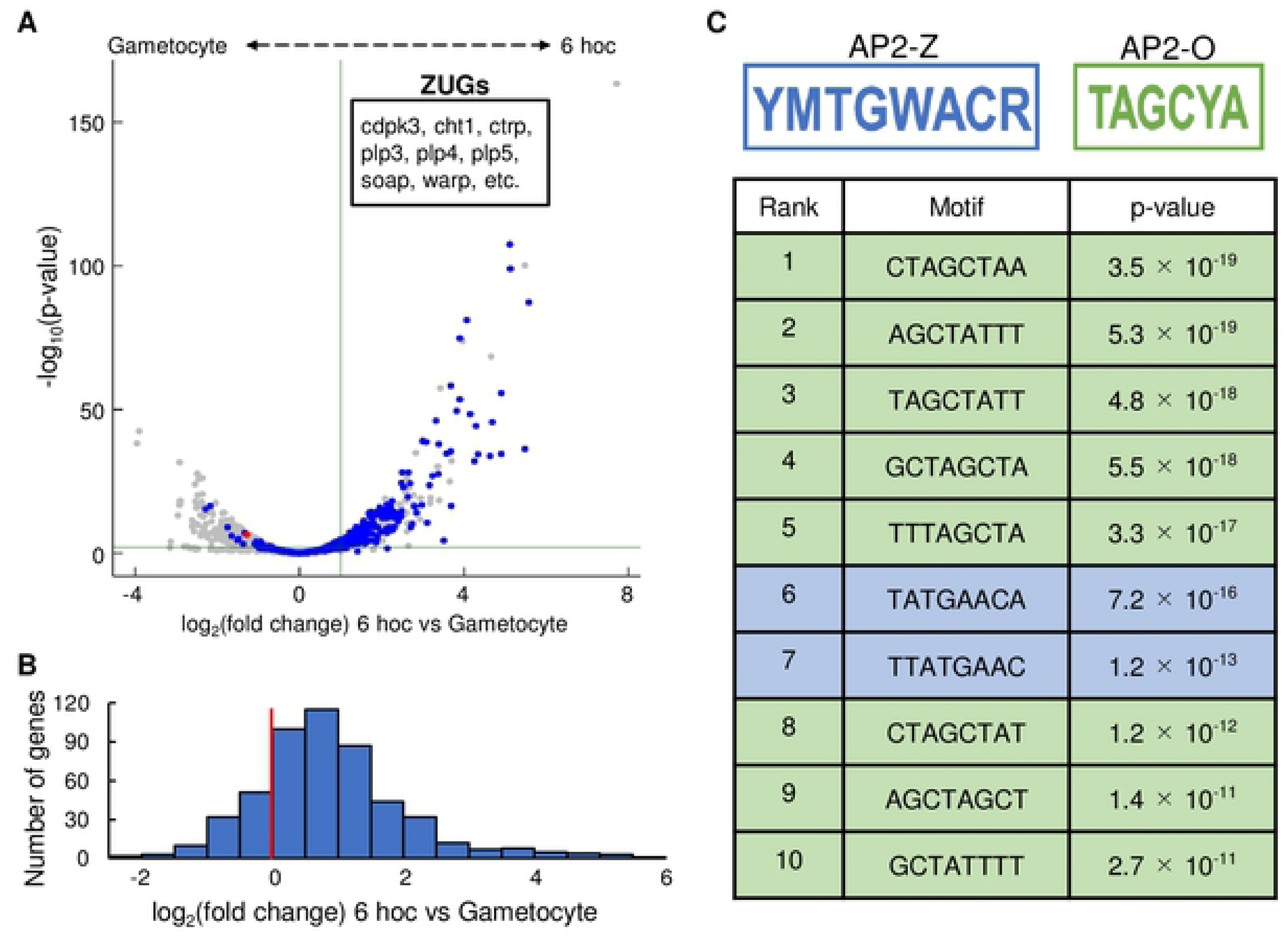
Differential expression analysis between gametocytes and zygotes at 6 hoc. (A) Volcano plot showing differential expression analysis between gametocytes and zygotes at 6 hoc. Red dot represents *ap2-z* and blue dots represent the target genes of AP2-Z. Green horizontal line indicates a *p*-value of 0.05 and a vertical line indicates a log_2_(fold change) of 1. (B) Histogram of the number of AP2-Z targets against the log_2_(fold change) value (bin = 0.5) in the differential expression analysis. Red vertical line indicates a log_2_(fold change) of 0. (C) Eight-base motifs enriched in the upstream of zygote- upregulated genes. AP2-Z motifs are indicated in blue and AP2-O motifs are indicated in green. Motifs were ranked according to their *p*-values.

To further confirm the relationship between the target genes of AP2-Z and the ZUGs, we investigated whether the binding motif of AP2-Z was enriched in the upstream region of the ZUGs. We assessed enrichment of any motif in the upstream region (300– 1,200 bp from the start codon) of ZUGs compared to that of the other genes by Fisher’s exact test. In the upstream region of ZUGs, TATGAACA, one of the binding motifs of AP2-Z [(T/C)(A/C)TG(A/T)AC(A/G)] was enriched, with a *p*-value of 7.2 × 10^-16^ (Fig 6C). Furthermore, another motif that partially contained the binding motif of AP2-Z, TTATGAAC, was also significantly enriched, with a *p*-value of 1.2 × 10^-13^. This result strongly indicated that AP2-Z functions as a major transcription factor during the early stage of zygote/ookinete development. The other enriched motifs, including the most enriched motif, fully or partially contained the binding motif of AP2-O [TAGC(T/C)A] (Fig 6C) [24]. Enrichment of the AP2-O binding motif could be due to the large overlap between the target genes of AP2-Z and AP2-O, which is evaluated later in detail (see Fig 8A). Notably, as with our result for the zygote stage, Witmer et al. reported a similar result for the mature ookinete stage, where the binding motif of AP2-O should be a major *cis*- element for upregulation of genes; in the upstream region of genes whose expression was upregulated more than 8-fold in mature ookinetes compared to female gametocytes, a motif almost identical to the AP2-Z binding motif, G(T/A)GAACA, was detected as the second most enriched motif (next to the binding motif of AP2-O) [32].

### Target genes of AP2-Z are downregulated in *ap2-z*(-)

To evaluate the contribution of AP2-Z to transcriptional regulation in zygotes, we performed differential expression analysis between WT and *ap2-z*(-) by RNA-seq. The analysis revealed that 161 genes, excluding *ap2-z*, were significantly downregulated (log_2_(fold change) < -1, *p*-value adj < 0.01) and 184 were significantly upregulated (log_2_(fold change) > 1, *p*-value adj < 0.01) in *ap2-z*(-) compared to WT at 6 hoc (Fig 7A and 7B, Table S4A and S4B). Of these downregulated genes, 86 genes were a target gene of AP2-Z, showing enrichment of the target genes in the downregulated genes (*p*-value = 1.8 × 10^-40^ by Fisher’s exact test). In addition, the mean value of log_2_(fold change) for the target genes was significantly lower than that for the other genes, with a *p*-value of 6.2 × 10^-65^ by Student’s t-test, indicating that the target genes were generally downregulated in *ap2-z*(-) (Fig 7B). Nevertheless, downregulation was only significant for a subset of the target genes (17%). We assumed that downregulation of some targets was not significant because they were already transcribed in female gametocytes, mostly by AP2-FG. Consistently, of 132 genes targeted by both AP2-Z and AP2-FG, only eight were significantly downregulated in *ap2-z*(-) (Fig 7C). In addition, the mean value of log_2_(fold change) for the common targets was significantly lower than that for the other AP2-Z targets, with a *p*-value of 4.8 × 10^-5^ by Student’s t-test.

**Fig 7.**
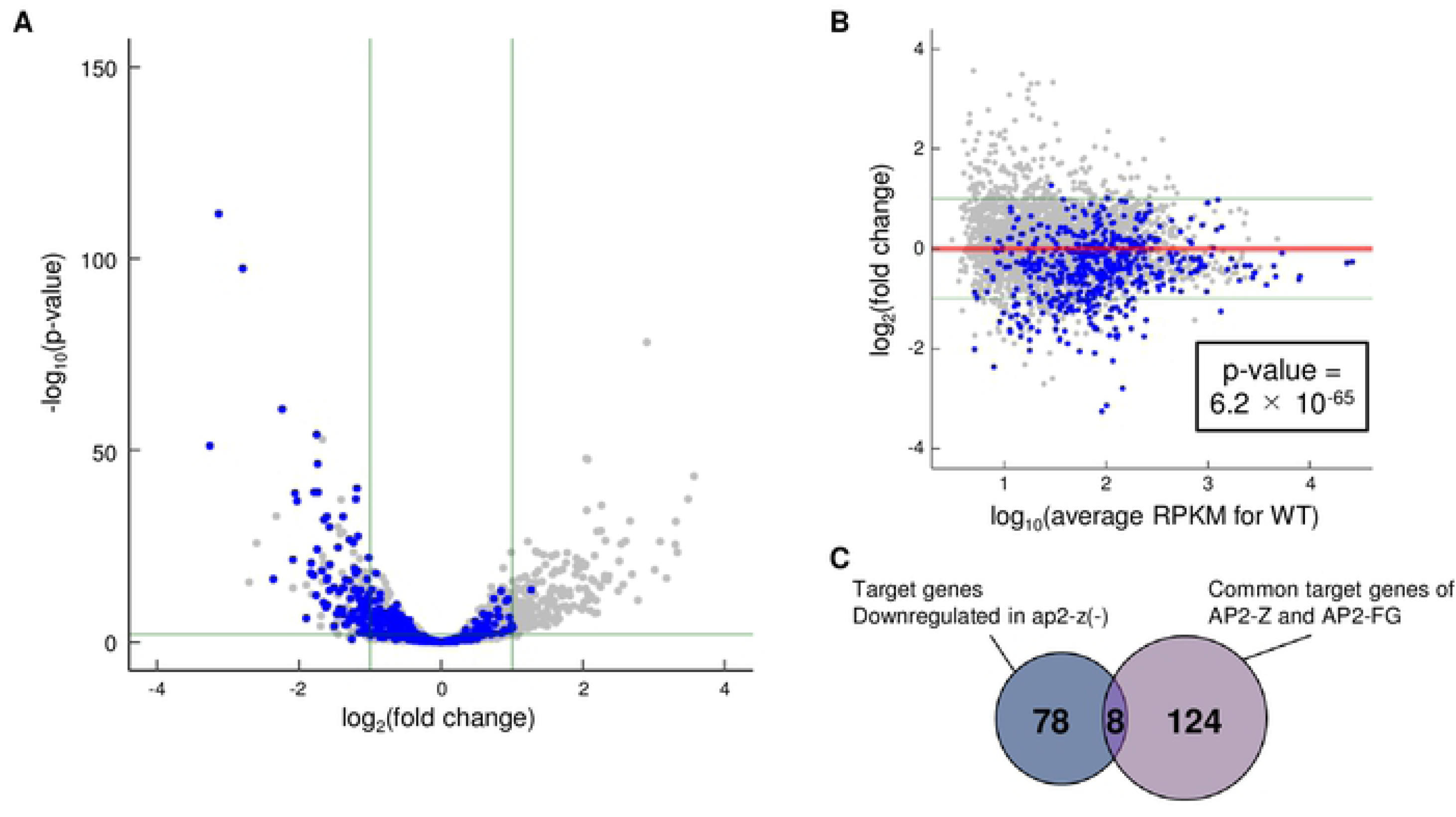
Differential expression analysis between WT and *ap2-z*(-) at 6 hoc. (A) Volcano plot showing differential expression analysis between WT and *ap2-z*(-) at 6 hoc. Blue dots represent the target genes of AP2-Z. Green horizontal line indicates a *p*- value of 0.05. Two vertical lines indicate log_2_(fold change) values of 1 and −1. The plot for *ap2-z* is omitted from the graph [log_2_(fold change) = −4.5, *p*-value adj = 4.9 × 10^-300^]. (B) MA plot showing differential expression analysis between WT and *ap2-z*(-) at 6 hoc. The *p*-value in the box represents a statistically significant difference between the mean values of log_2_(fold change) for AP2-Z targets and the other genes. Two green lines indicate log_2_(fold change) values of 1 and −1, and the red line indicates a log_2_(fold change) of 0. (C) Venn diagram showing the overlap between AP2-Z targets downregulated in *ap2-z*(-) and the common targets of AP2-Z and AP2-FG.

### Comparative targetome analyses between AP2-FG, AP2-Z, and AP2-O suggests their roles in regulating zygote/ookinete development

The results of this study suggested that gene expression during *Plasmodium* zygote/ookinete development is regulated by AP2-FG, as female-stored mRNA, and AP2- Z and AP2-O, as *de novo* transcripts. In this regard, we attempted to investigate the role of each of these three transcription factors by comparing their target genes.

First, we compared the target genes of AP2-Z and AP2-O. The two target gene sets showed a large overlap (307 genes), accounting for 60% of the targets of AP2-Z (Fig 8A, Table S2). This result was consistent with the above result that the binding motif of AP2-O was enriched in the upstream of ZUGs. These overlapped genes included most of the known microneme genes, ookinete surface protein genes, and IMC/pellicle genes, which are all related to parasite invasion. Therefore, the main role of AP2-Z may be promoting the development of an invasive stage until expression of AP2-O. Accordingly, these common targets are continuously activated by the two transcription factors throughout development from zygote to ookinete, which may be important for efficiently promoting ookinete formation. These common target genes have ChIP binding sites of both AP2-Z and AP2-O in their upstream region. Notably, these binding sites of AP2-Z and AP2-O are located very close to each other in the upstream region of the same genes (mostly within 100 bp). (Fig 8B and 8C). This result suggested that the two transcription factors share the same regulatory regions for their common target genes, and that a major transcriptional regulator gradually switches from AP2-Z to AP2-O in these regulatory regions as zygote/ookinete development progresses.

**Fig 8.**
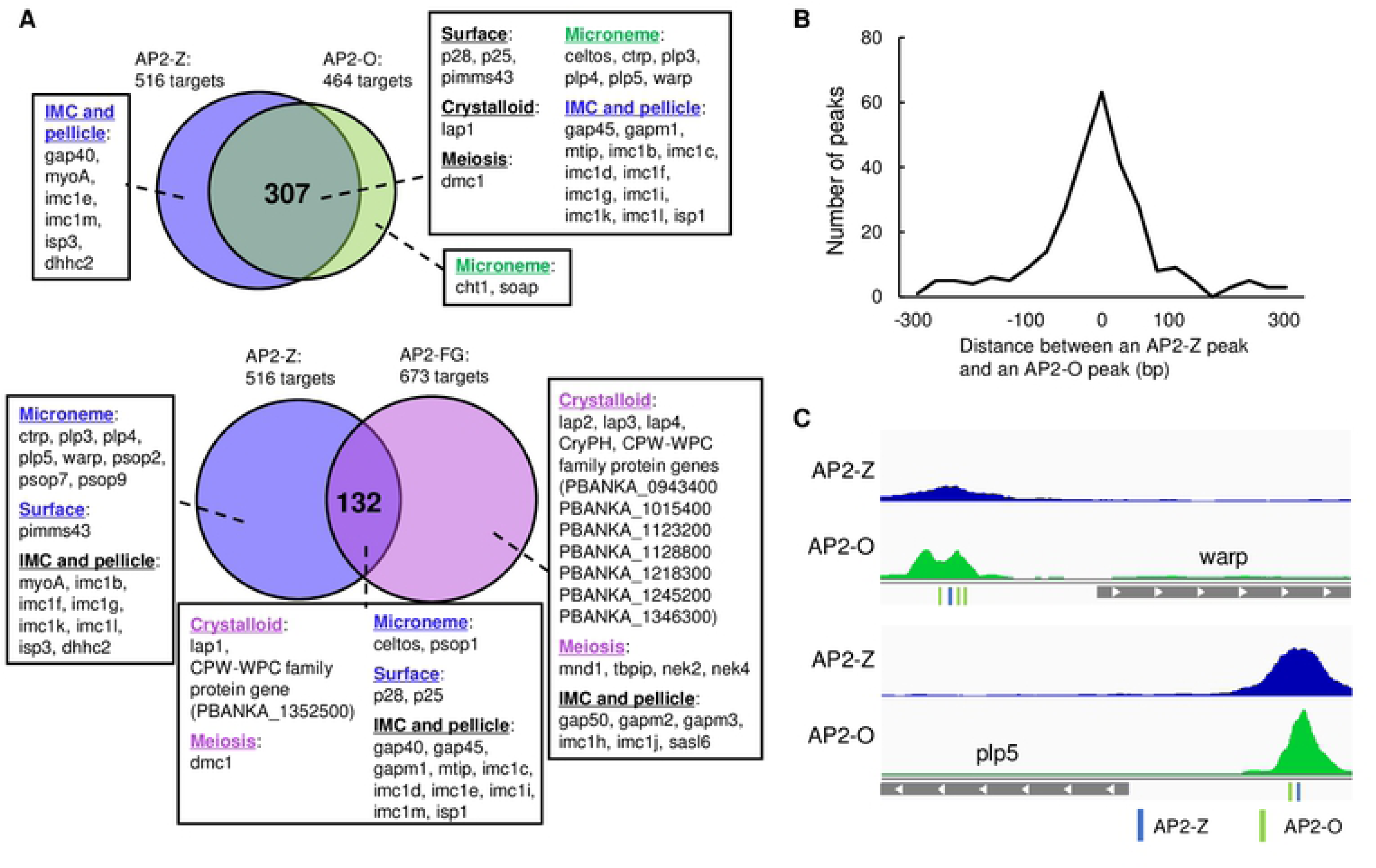
Comparison of the target genes between AP2-Z, AP2-O, and AP2-FG. (A) Venn diagrams showing the overlap between the target genes of AP2-Z and AP2-O (top) and AP2-Z and AP2-FG (bottom). Genes were assigned according to previous literature [24,34–42,48–73]. (B) Distance between ChIP peaks of AP2-Z and AP2-O in the upstream region of their common targets. (C) Representative peak images of the motifs are indicated in blue and green, respectively.

In contrast, the target genes unique to AP2-Z or AP2-O contained only a small number of known ookinete genes. These ookinete genes in the unique targets contained six known IMC/pellicle genes for AP2-Z and two known microneme genes, *cht1* and *soap*, for AP2-O (Fig 8A, Table S2). While IMC formation is necessary from the first steps of ookinete formation, *i.e.* determination of apical polarity and construction of an apical protrusion [33], microneme is essential for parasite motility and invasion in mature ookinetes. Therefore, these results might suggest that, while regulating the common targets, AP2-Z and AP2-O also regulate their unique target genes in accordance with their own expression pattern during zygote/ookinete development.

Next, we investigated the target genes of AP2-Z and AP2-FG. Since the target genes of AP2-FG contains most of the female-enriched genes [13], we considered that this analysis could possibly provide an indication for the role of female-stored mRNAs unique from that of *de novo* transcripts. The genes common in the two target sets contained 132 genes, representing approximately 25% of the AP2-Z targets (Fig 8A, Table S2). Several IMC/pellicle genes were included in the target genes of AP2-FG as both common and unique targets (Fig 8A, Table S2), which, as with previous studies [13,18,19], indicated that female-stored genes are essential for ookinete formation. This is consistent with the fact that fertilized females were able to develop into retort-form ookinetes, even when *ap2-z* was disrupted [10] or fertilized females were treated with a transcriptional inhibitor [19]. On the other hand, the AP2-FG targets contained only two known microneme genes, both of them as common targets, suggesting that although playing a role in the initial steps of ookinete formation, female-stored mRNAs may not largely contribute to the later steps. The other unique targets of AP2-FG contained most of the genes related to meiosis and crystalloid biogenesis (Fig 8A, Table S2), which are not directly involved in ookinete formation [34–38]. This result suggested that developmental processes other than ookinete formation are mainly regulated by female-stored mRNAs during *Plasmodium* zygote/ookinete development.

### Unique targets of AP2-Z contain an important gene for ookinete development

Although the target genes of AP2-Z largely overlapped with those of AP2-O, they also contained a considerable number of unique targets, most of which have not yet been functionally assessed. To evaluate the roles of these unique targets, we assessed one of the unique target genes with an unknown function, PBANKA_0508300 (hereafter named ookinete maturation gene 1 (*omg1*)). Notably, *omg1* was significantly upregulated at 6 hoc compared to gametocytes, with a log_2_(fold change) of 3.33, suggesting its transcription after fertilization. By fusing *mNG* to *omg1* using CRISPR/Cas9 (OMG1::mNG) (Fig S1C), we investigated the *omg1* expression pattern during ookinete development. In the blood stage, both female and male gametocytes of OMG1::mNG showed no fluorescent signal (Fig 9A). After fertilization, a weak signal started to appear at 6 hoc. This signal intensified in retort-form ookinetes and was still observed in matured ookinetes. Fluorescence was observed along the periphery of ookinetes but was absent in the zygote remnant and the apical end (Fig 9A). This pattern of distribution is similar to that of several known IMC components, such as IMC1h and IMC1i [24,39–42]. Therefore, this result suggested that OMG1 is a component of IMC that plays a role in pellicle formation. This was consistent with the comparative targetome analysis between AP2-Z and AP2-O, in which it was suggested that unique targets of AP2-Z include genes required from the first steps of ookinete formation, such as IMC/pellicle genes.

**Fig 9.**
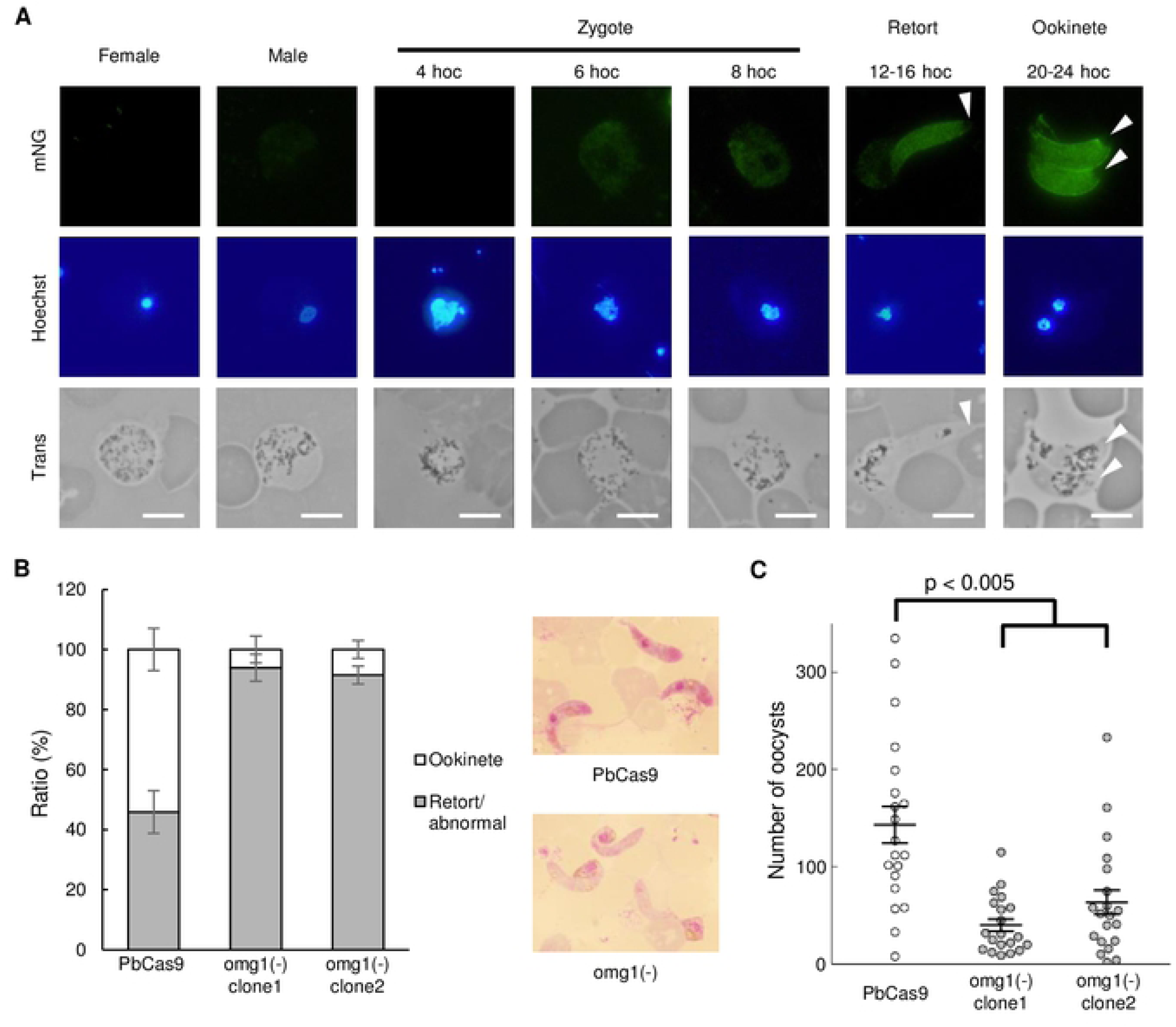
Investigation of one of the unique targets of AP2-Z, *omg1*. (A) Expression of OMG1 in the OMG1::mNG parasite during sexual development. Arrow heads indicate the apical end of the ookinete. Nuclei were stained with Hoechst 33342. Scale bar = 5 μm. (B) Ratio of ookinetes and retort-form/abnormal zygotes to all fertilized cells in PbCas9 and *omg1*(-) at 16 hoc. Right side of the figure shows Giemsa-stained images of ookinetes at 16 hoc in PbCas9 (top) and *omg1*(-) (bottom). Error bars indicate the standard error of the mean (n = 3). (C) Number of midgut oocysts at 14 days post infection for PbCas9 and *omg1*(-). Lines indicate the mean values and the standard error (n=20). The *p*-value on the graph was calculated by Student’s t-test.

We next disrupted *omg1* by CRISPR/Cas9 [*omg1*(-)] using the PbCas9 parasite, obtaining two clonal lines from two independent transfection experiments (Fig S1D), and investigated its phenotype. Both female and male gametocytes of *omg1*(-) appeared normal and were able to fertilize. However, ookinete maturation was delayed in *omg1*(-) compared to PbCas9 parasites. In *omg1*(-), the ratio of banana-shaped ookinetes to all fertilized cells was less than 10% at 16 hoc for both clones, whereas more than half of fertilized cells had become banana-shaped ookinetes in PbCas9 at the same time point (Fig 9B). Nonetheless, disruption of *omg1* did not completely impair ookinete development as most fertilized cells became banana-shaped ookinetes prior to 30 hoc.

We further investigated whether the delay in ookinete maturation affects mosquito infectivity. Mice infected with PbCas9 parasites or *omg1*(-) were fed on *Anopheles stephensi* mosquitoes. At 14 days post infection, the number of oocysts in the midgut of infected mosquitoes was significantly reduced in both clones of *omg1*(-) compared to in PbCas9 (Fig 9C). Therefore, because of impaired ookinete development, *omg1*(-) might delay in penetrating the peritrophic membrane to reach the basal lamina and hence be exposed to the hostile environment of the mosquito midgut for a longer time. Collectively, these results suggested that the unique targets of AP2-Z contain genes that are important for ookinete maturation.

## Discussion

Previous studies showed that, in *P. berghei*, female-stored mRNA is essential for zygote/ookinete development; thus, disruption of the DOZI complex completely abolishes meiosis and the ookinete conversion of fertilized females [15, 18]. Moreover, fertilized females are able to develop until the retort stage of ookinetes in the presence of a transcription inhibitor, which suggested that female-stored mRNAs are sufficient for promoting the initial step of ookinete conversion [19]. On the other hand, our previous studies revealed the essential role of *de novo* transcription by AP2-O in ookinete maturation, where disruption of *ap2-o* resulted in developmental arrest at the retort stage of ookinetes [23]. Putting these studies together suggests that *Plasmodium* zygote/ookinete development is promoted by female-stored mRNA until the retort stage of ookinetes and *de novo* transcription by AP2-O in the later stage. Accordingly, transcriptional activity in zygotes has been considered quiescent, similar to animal embryos that are transcriptionally silent in the early stage after fertilization [21, 22]. In this study, by investigating the roles of a novel AP2-family transcription factor, AP2-Z, we demonstrated that *de novo* transcription in *P. berghei* zygotes is actually activated by AP2-Z at the zygote stage and is essential for zygote/ookinete development. Therefore, our results indicated that female-stored mRNAs and *de novo* transcripts by AP2-Z both promote the early stages of zygote/ookinete development.

Transcriptional regulation in Plasmodium is very simple; a single DNA-binding transcription factor, such as AP2-G and AP2-O, comprehensively regulates certain stage-specific genes [10,13,24,43,44]. Such a simple mechanism may enable regulation of the entire *Plasmodium* life cycle by only a small number of sequence-specific transcription factors (approximately 30 AP2-family transcription factors and a few others have been identified to date [45–47]). If the life cycle is simply proceeded by sequential expression of such master transcription factors, it is plausible to hypothesize that they make up a large cascade covering the entire life cycle; i.e., every stage-specific transcription factor, while regulating the respective stage-specific genes, activates the transcription factors responsible for regulation of the next stage. In this study, we confirmed such a cascade during sexual development via the promoter assay of ap2-z; that is, transcription of ap2-z in female gametocytes was completely depleted upon disruption of the binding motifs of AP2-G and AP2-FG in the upstream region. Furthermore, through target analysis of AP2-Z, we revealed that AP2-Z targets ap2-o and ap2-o2, which are essential transcription factors for ookinete maturation. These results indicated that a dynamic cascade of transcription factors that starts from AP2-G occurs during the entire process of Plasmodium sexual development (Fig 10A and 10B). If such a simple cascade of transcription factors was found throughout the entire life cycle of Plasmodium, we should always be able to identify transcriptional regulators for the next developmental stage from target analysis of a certain transcriptional activator, just as we found ap2-z from the target genes of AP2-G and AP2-FG in this study.

**Fig 10.**
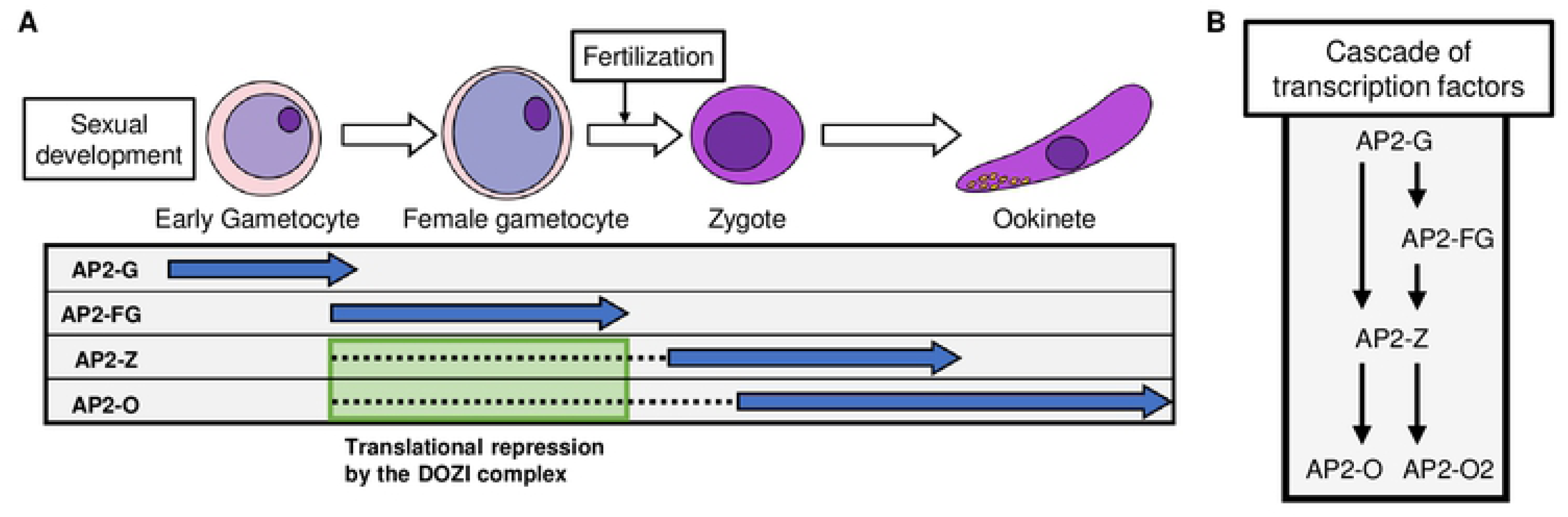
Cascade of transcription factors during the sexual development of *Plasmodium*. (A) Schematic illustration of the sequential expression of sexual transcription factors: AP2-G, AP2-FG, AP2-Z, and AP2-O. Blue arrows indicate a period of protein expression and dashed lines indicate a period of translational repression. (B) Schematic illustration showing the hierarchical relationship of sexual transcription factors during *Plasmodium* sexual development.

In conclusion, this study revealed the essential role of transcription by AP2-Z in zygotes and a dynamic cascade of transcription factors during *Plasmodium* sexual development. Moreover, our comparative targetome analyses provided new insights into the mechanisms regulating zygote/ookinete development. To further expand our knowledge of the molecular mechanisms underlying zygote/ookinete development and transmission from vertebrate hosts to mosquitoes, it is important to evaluate the functions of individual target genes. Especially, we believe that investigation of the unique target genes of AP2-Z could help us elucidate new aspects of the mechanism regulating ookinete development because these genes are not readily identified in transcriptome analyses of gametocytes or ookinetes. In addition, although most of the unique target genes of AP2-Z have not yet been functionally investigated, they may well be important for zygote/ookinete development, as shown in this study with *omg1*.

## Materials and Methods

### Ethical statement

All experiments in this study were performed according to recommendations in the Guide for the Care and Use of Laboratory Animals of the National Institutes of Health in order to minimize animal suffering. All protocols were approved by the Animal Research Ethics Committee of Mie University (permit number 23–29).

### Parasite preparation

All parasites used in this study were inoculated in Balb/c or ddY mice. The *ap2-z*(-) and DOZI::GFP parasites were derived from the WT ANKA strain, and all other transgenic parasites were generated by the CRISPR/Cas9 system using Cas9-expressing parasites called PbCas9 [28]. Ookinete cultures were performed as follows. First, infected blood was intraperitoneally injected into phenylhydrazine (nacalai tesque)-treated mice. When the parasitemia reached approximately 3%, the mice were fed sulfadiazine (Sigma) in their drinking water (10 μg/ml) to kill asexual stage parasites. After two days of sulfadiazine treatment, infected blood was withdrawn from the mice and diluted in culture medium (RPMI1640 medium whose pH was adjusted to 8.0 with NaOH solution, supplemented with fetal calf serum and penicillin/streptomycin, achieving final concentrations of 20% and 1%, respectively). Blood samples were then passed through a column filled with cellulose powder CF11 or a Plasmodipur filter for ChIP-seq and RNA-seq experiments, and the parasites were incubated at 20 °C. Midgut oocysts and sporozoites were counted from the midgut of infected mosquitoes at 14 days post-infective blood meal.

### Generation of transgenic parasites

For tagging DOZI with GFP, the conventional homologous recombination method was used as previously reported [23]. Briefly, two homologous regions were cloned into the *gfp*-fusion vector to fuse *dozi* in frame with *gfp*. The vector was linearized by restriction enzymes and transfected into parasites by electroporation. Mutants were selected with 70 μg/mL pyrimethamine in drinking water. The previously reported CRISPR/Cas9 system was used to generate transgenic parasites by CRISPR/Cas9 [28]. Briefly, PbCas9 parasites, which constitutively express Cas9 endonuclease, were transfected with donor DNA prepared by the overlap PCR method and sgRNA vector by electroporation. After transfection, mice infected with the transfectants were treated with 70 μg/mL of pyrimethamine (Sigma) in their drinking water for three days to select the desired mutants. All clonal parasites were obtained by limiting dilution methods, and genotyping was performed by PCR and Sanger sequencing. All primers used in this study are listed in Table S5 (No. 1–39).

### RNA immunoprecipitation experiments

For the RIP experiments, mice infected with DOZI::GFP parasites were treated with sulfadiazine in their drinking water to kill asexual parasites. After two days of drug treatment, whole blood was withdrawn from the infected mice and washed once with RPMI medium. After lysing red blood cells in ice-cold lysis solution (1.5 M NH4Cl, 0.1 M KHCO_3_, 10 mM EDTA), the samples were subjected to RIP experiments, which were performed using RiboCluster Profiler (MBL Co., Ltd.) according to the manufacturer’s instructions. Briefly, cells were lysed in the provided lysis buffer, and the lysate was mixed with antibody-free Protein A agarose beads then incubated at 4 °C as a preclear step. Next, the precleared cell lysate was separated from the beads by centrifugation. A small portion of the precleared sample was collected for input RNA quantification, and the rest was mixed with antibody-immobilized Protein A agarose beads and incubated at 4 °C. During this process, anti-GFP polyclonal antibody (Abcam, ab290) and normal rabbit IgG supplied in the kit were used for immunoprecipitation of DOZI-associated RNAs and as a negative control, respectively. Finally, RNA was purified from the beads and subjected to RT-qPCR analysis. Three biologically independent samples were prepared and used for the analysis.

### RT-qPCR analysis

cDNA was synthesized from total RNA or immunoprecipitated RNA using PrimeScript RT reagent Kit with gDNA Eraser (Takara). RT-qPCR analysis was performed using TB Green Fast qPCR Mix (Takara) and Thermal Cycler Dice Real Time System II (Takara). All primers used in this study are listed in Table S5 (No. 40–45).

### ChIP-seq and sequencing data analysis

The ChIP-seq experiments were performed as described previously [24]. Briefly, AP2- Z::GFP parasites at 6 hoc were fixed by adding formalin solution to achieve the final concentration of 1% and incubating the solution at 30 °C. After fixing, red blood cells were lysed in ice-cold 1.5 M NH4Cl solution. Next, the residual cells were lysed in SDS lysis buffer, and the lysate was sonicated using Bioruptor (Cosmo Bio) to shear chromatin. A small portion of the sonicated lysate was collected as input samples. ChIP samples were immunoprecipitated from the sonicated lysate with anti-GFP polyclonal antibodies (Abcam, ab290) immobilized on Dynabeads Protein A (Invitrogen). DNA fragments were purified from the ChIP and input samples, and then used for library construction, which was performed with a KAPA HyperPrep Kit according to the manufacturer’s instructions. The library was then sequenced by Illumina NextSeq. Two biologically independent experiments were performed.

The obtained sequence data were mapped onto the reference genome sequence of *P. berghei* ANKA, downloaded from PlasmoDB 46, by Bowtie2. Reads that aligned more than two times were removed from the mapping data. Using the mapping data from the ChIP and input samples, peaks were identified using the macs2 callpeak function with fold enrichment > 4.0 and *q*-value < 0.01, and common peaks between the two experiments were used for further analysis. Prediction of binding motifs was performed by the Fisher’s exact test. Genes that had peaks within the region upstream of 1,200 bp from ATG were identified as target genes. Parameters for all programs were set to the default, unless specified otherwise.

### RNA-seq and sequence data analysis

Total RNA was extracted from Plasmodipur-filtered mouse blood infected with WT or *ap2-z*(-) using the Isogen II reagent (Nippon gene). Briefly, red blood cells were lysed in ice-cold 1.5 M NH_4_Cl solution. The residual cells were then lysed in Isogen II, and total RNA was purified from the lysate according to the manufacturer’s instructions. From the total RNA, RNA-seq libraries were prepared using the KAPA mRNA HyperPrep Kit and sequenced by Illumina NextSeq. Three biologically independent experiments were performed for each sample. The obtained sequence data were mapped onto the reference genome sequence of *P. berghei* by HISAT2, setting the maximum intron length threshold to 1,000. The mapping data for each sample were analyzed by featureCounts and compared using DESeq2. The reads per kilobase of transcript per million mapped reads for each gene were calculated from the featureCounts results; genes with values of less than five in all three datasets for WT at 6 hoc were removed from the differential expression analysis. Genes in subtelomeric regions were also removed. For all programs, the parameters were set to the default, unless specified otherwise.

## Supporting information

**Fig S1.**
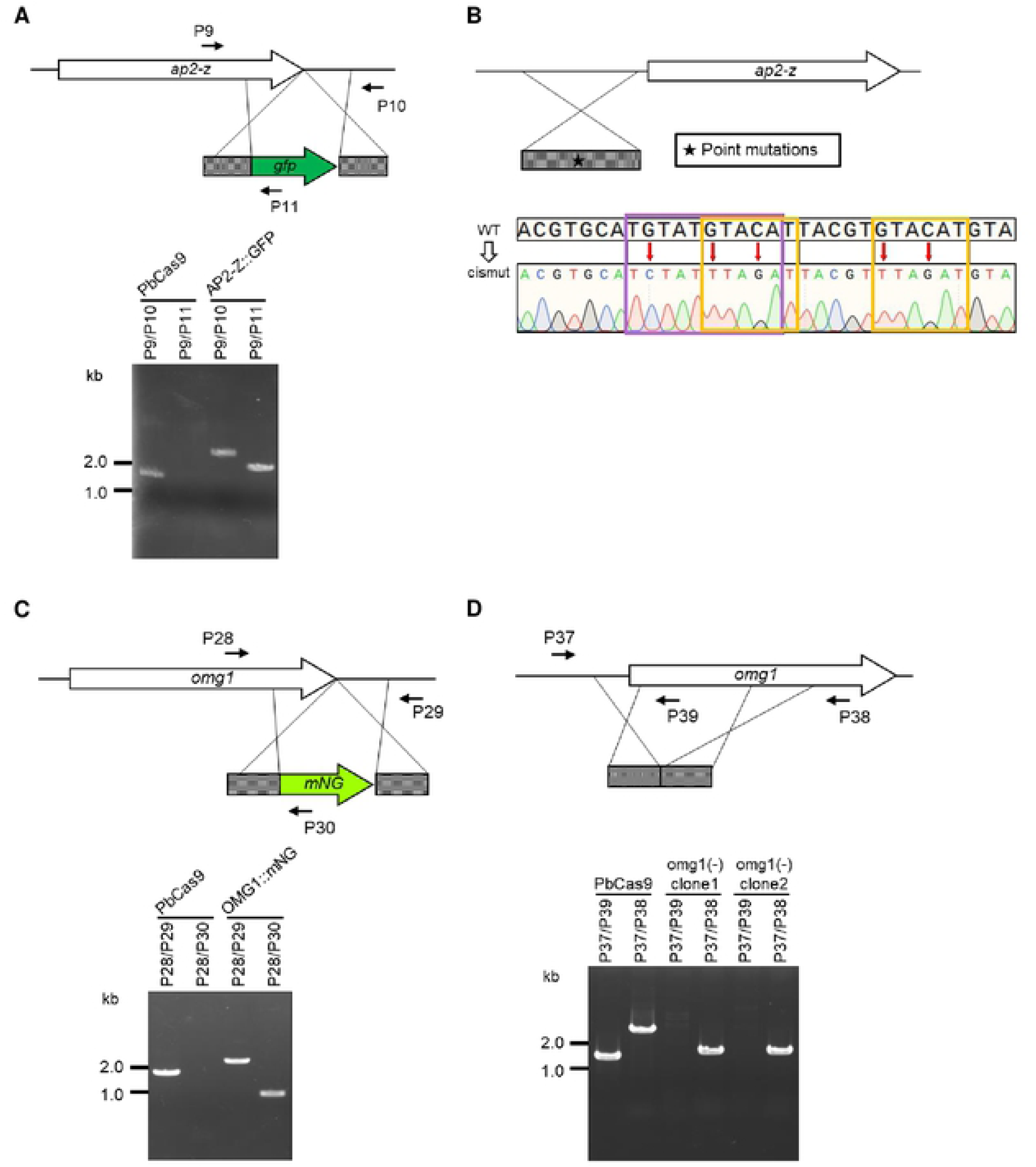
Genotyping of transgenic parasites developed in this study. (A) AP2-Z::GFP. (B) *ap2-z*(-). (C) OMG1::mNG. (D) *omg1*(-).

**Table S1. List of peaks identified in the ChIP-seq experiments.** (A) Experiment 1. (B) Experiment 2.

**Table S2. List of target genes of AP2-Z.**

**Table S3. List of zygote-upregulated genes.**

**Table S4. List of differentially expressed genes in *ap2-z*(-).** (A) Significantly downregulated genes. (B) Significantly upregulated genes.

**Table S5. List of primers used in this study.**

## References

1. World Health Organization. World Malaria Report. World Health. 2021. doi:ISBN 978 92 4 1564403

2. Bousema T, Drakeley C. Epidemiology and infectivity of Plasmodium falciparum and Plasmodium vivax gametocytes in relation to malaria control and elimination. Clinical Microbiology Reviews. American Society for Microbiology 1752 N St., N.W., Washington, DC; 2011. pp. 377–410. doi:10.1128/CMR.00051-10

3. Guttery DS, Holder AA, Tewari R. Sexual development in plasmodium: Lessons from functional analyses. PLoS Pathog. 2012;8: e1002404. doi:10.1371/journal.ppat.1002404

4. Baker DA. Malaria gametocytogenesis. Molecular and Biochemical Parasitology. Elsevier; 2010. pp. 57–65. doi:10.1016/j.molbiopara.2010.03.019

5. Josling GA, Llinás M. Sexual development in Plasmodium parasites: Knowing when it’s time to commit. Nature Reviews Microbiology. Nature Publishing Group; 2015. pp. 573–587. doi:10.1038/nrmicro3519

6. Shahabuddin M. Plasmodium ookinete development in the mosquito midgut: A case of reciprocal manipulation. Parasitology. 1998;116. doi:10.1017/s0031182000084973

7. Sinha A, Hughes KR, Modrzynska KK, Otto TD, Pfander C, Dickens NJ, et al. A cascade of DNA-binding proteins for sexual commitment and development in Plasmodium. Nature. 2014;507: 253–257. doi:10.1038/nature12970

8. Kafsack BFC, Rovira-Graells N, Clark TG, Bancells C, Crowley VM, Campino SG, et al. A transcriptional switch underlies commitment to sexual development in malaria parasites. Nature. 2014;507: 248–252. doi:10.1038/nature12920

9. Kent RS, Modrzynska KK, Cameron R, Philip N, Billker O, Waters AP. Inducible developmental reprogramming redefines commitment to sexual development in the malaria parasite Plasmodium berghei. Nat Microbiol. 2018;3: 1206–1213. doi:10.1038/s41564-018-0223-6

10. Yuda M, Kaneko I, Murata Y, Iwanaga S, Nishi T. Mechanisms of triggering malaria gametocytogenesis by AP2-G. Parasitol Int. 2021;84: 102403. doi:10.1016/j.parint.2021.102403

11. Yuda M, Iwanaga S, Kaneko I, Kato T. Global transcriptional repression: An initial and essential step for Plasmodium sexual development. Proc Natl Acad Sci U S A. 2015;112: 12824–12829. doi:10.1073/pnas.1504389112

12. Singh S, Santos JM, Orchard LM, Yamada N, van Biljon R, Painter HJ, et al. The PfAP2-G2 transcription factor is a critical regulator of gametocyte maturation. Mol Microbiol. 2021;115: 1005–1024. doi:10.1111/MMI.14676

13. Yuda M, Kaneko I, Iwanaga S, Murata Y, Kato T. Female-specific gene regulation in malaria parasites by an AP2-family transcription factor. Mol Microbiol. 2020;113: 40–51. doi:10.1111/mmi.14334

14. Li Z, Cui H, Guan J, Liu C, Yang Z, Yuan J. Plasmodium transcription repressor AP2-O3 regulates sex-specific identity of gene expression in female gametocytes. EMBO Rep. 2021;22: 1–18. doi:10.15252/embr.202051660

15. Mair GR, Braks JAM, Garver LS, Wiegant JCAG, Hall N, Dirks RW, et al. Regulation of sexual development of Plasmodium by translational repression. Science (80- ). 2006;313: 667–669. doi:10.1126/science.1125129

16. Mair GR, Braks JAM, Garver LS, Dimopoulos G, Hall N, Wiegant JGAG, et al. Translational Repression is essential for Plasmodium sexual development and mediated by a DDX6-type RNA helicase. Science (80- ). 2006;313: 667–669. Available: http://www.ncbi.nlm.nih.gov/pmc/articles/PMC1609190/

17. Braks JAM, Mair GR, Franke-Fayard B, Janse CJ, Waters AP. A conserved U- rich RNA region implicated in regulation of translation in Plasmodium female gametocytes. Nucleic Acids Res. 2008;36: 1176–1186. doi:10.1093/nar/gkm1142

18. Mair GR, Lasonder E, Garver LS, Franke-Fayard BMD, Carret CK, Wiegant JCAG, et al. Universal features of post-transcriptional gene regulation are critical for Plasmodium zygote development. PLoS Pathog. 2010;6. doi:10.1371/journal.ppat.1000767

19. Guerreiro A, Deligianni E, Santos JM, Silva PAGC, Louis C, Pain A, et al. Genome-wide RIP-Chip analysis of translational repressor-bound mRNAs in the Plasmodium gametocyte. Genome Biol. 2014;15: 1–16. doi:10.1186/s13059-014-0493-0

20. Waters AP, Van Spaendonkth RML, Ramesar J, Vervenne RAW, Dirksll RW, Thompsont J, et al. Species-specific regulation and switching of transcription between stage-specific ribosomal RNA genes in Plasmodium berghei. J Biol Chem. 1997;272: 3583–3589. doi:10.1074/jbc.272.6.3583

21. Baroux C, Autran D, Gillmor CS, Grimanelli D, Grossniklaus U. The maternal to zygotic transition in animals and plants. Cold Spring Harbor Symposia on Quantitative Biology. Cold Spring Harbor Laboratory Press; 2008. pp. 89–100. doi:10.1101/sqb.2008.73.053

22. Lee MT, Bonneau AR, Giraldez AJ. Zygotic genome activation during the maternal-to-zygotic transition. Annu Rev Cell Dev Biol. 2014;30: 581–613. doi:10.1146/annurev-cellbio-100913-013027

23. Yuda M, Iwanaga S, Shigenobu S, Mair GR, Janse CJ, Waters AP, et al. Identification of a transcription factor in the mosquito-invasive stage of malaria parasites. Mol Microbiol. 2009;71: 1402–1414. doi:10.1111/j.1365-2958.2009.06609.x

24. Kaneko I, Iwanaga S, Kato T, Kobayashi I, Yuda M. Genome-Wide Identification of the Target Genes of AP2-O, a Plasmodium AP2-Family Transcription Factor. PLoS Pathog. 2015;11: 1–27. doi:10.1371/journal.ppat.1004905

25. Modrzynska K, Pfander C, Chappell L, Yu L, Suarez C, Dundas K, et al. A Knockout Screen of ApiAP2 Genes Reveals Networks of Interacting Transcriptional Regulators Controlling the Plasmodium Life Cycle. Cell Host Microbe. 2017;21: 11–22. doi:10.1016/j.chom.2016.12.003

26. Zhang C, Li Z, Cui H, Jiang Y, Yang Z, Wang X, et al. Systematic CRISPR-Cas9-mediated modifications of plasmodium yoelii ApiAP2 genes reveal functional insights into parasite development. MBio. 2017;8: 1–17. doi:10.1128/mBio.01986-17

27. Balaji S, Madan Babu M, Iyer LM, Aravind L. Discovery of the principal specific transcription factors of Apicomplexa and their implication for the evolution of the AP2-integrase DNA binding domains. Nucleic Acids Res. 2005;33: 3994–4006. doi:10.1093/nar/gki709

28. Shinzawa N, Nishi T, Hiyoshi F, Motooka D, Yuda M, Iwanaga S. Improvement of CRISPR/Cas9 system by transfecting Cas9-expressing Plasmodium berghei with linear donor template. Commun Biol 2020 31. 2020;3: 1–13. doi:10.1038/s42003-020-01138-2

29. van Dijk MR, van Schaijk BCL, Khan SM, van Dooren MW, Ramesar J, Kaczanowski S, et al. Three members of the 6-cys protein family of plasmodium play a role in gamete fertility. PLoS Pathog. 2010;6: 1–13. doi:10.1371/journal.ppat.1000853

30. Bateman E. Autoregulation of eukaryotic transcription factors. Progress in nucleic acid research and molecular biology. Academic Press; 1998. pp. 133– 168. doi:10.1016/s0079-6603(08)60892-2

31. Crews ST, Pearson JC. Transcriptional autoregulation in development. Current Biology. Cell Press; 2009. pp. R241–R246. doi:10.1016/j.cub.2009.01.015

32. Witmer K, Fraschka SA, Vlachou D, Bártfai R, Christophides GK. An epigenetic map of malaria parasite development from host to vector. Sci Rep. 2020;10: 1– 19. doi:10.1038/s41598-020-63121-5

33. Ferreira JL, Heincke D, Wichers JS, Liffner B, Wilson DW, Gilberger TW. The Dynamic Roles of the Inner Membrane Complex in the Multiple Stages of the Malaria Parasite. Frontiers in Cellular and Infection Microbiology. Frontiers Media S.A.; 2021. p. 841. doi:10.3389/fcimb.2020.611801

34. Mlambo G, Coppens I, Kumar N. Aberrant Sporogonic Development of Dmc1 (a Meiotic Recombinase) Deficient Plasmodium berghei Parasites. PLoS One. 2012;7: e52480. doi:10.1371/journal.pone.0052480

35. Pradel G, Hayton K, Aravind L, Iyer LM, Abrahamsen MS, Bonawitz A, et al. A multidomain adhesion protein family expressed in Plasmodium falciparum is essential for transmission to the mosquito. J Exp Med. 2004;199: 1533–1544. doi:10.1084/jem.20031274

36. Carter V, Shimizu S, Arai M, Dessens JT. PbSR is synthesized in macrogametocytes and involved in formation of the malaria crystalloids. Mol Microbiol. 2008;68: 1560–1569. doi:10.1111/j.1365-2958.2008.06254.x

37. Lavazec C, Moreira CK, Mair GR, Waters AP, Janse CJ, Templeton TJ. Analysis of mutant Plasmodium berghei parasites lacking expression of multiple PbCCp genes. Mol Biochem Parasitol. 2009;163: 1–7. doi:10.1016/j.molbiopara.2008.09.002

38. Saeed S, Tremp AZ, Dessens JT. Biogenesis of the crystalloid organelle in Plasmodium involves microtubule-dependent vesicle transport and assembly. Int J Parasitol. 2015;45: 537–547. doi:10.1016/j.ijpara.2015.03.002

39. Volkmann K, Pfander C, Burstroem C, Ahras M, Goulding D, Rayner JC, et al. The alveolin IMC1h is required for normal ookinete and sporozoite motility behaviour and host colonisation in plasmodium berghei. PLoS One. 2012;7: e41409. doi:10.1371/journal.pone.0041409

40. Tremp AZ, Al-Khattaf FS, Dessens JT. Distinct temporal recruitment of Plasmodium alveolins to the subpellicular network. Parasitol Res. 2014;113: 4177–4188. doi:10.1007/s00436-014-4093-4

41. Al-Khattaf FS, Tremp AZ, Dessens JT. Plasmodium alveolins possess distinct but structurally and functionally related multi-repeat domains. Parasitol Res. 2015;114: 631–639. doi:10.1007/s00436-014-4226-9

42. Gao H, Yang Z, Wang X, Qian P, Hong R, Chen X, et al. ISP1-Anchored Polarization of GCβ/CDC50A Complex Initiates Malaria Ookinete Gliding Motility. Curr Biol. 2018;28: 2763–2776.e6. doi:10.1016/j.cub.2018.06.069

43. Santos JM, Josling G, Ross P, Joshi P, Orchard L, Campbell T, et al. Red Blood Cell Invasion by the Malaria Parasite Is Coordinated by the PfAP2-I Transcription Factor. Cell Host Microbe. 2017;21: 731–741.e10. doi:10.1016/J.CHOM.2017.05.006

44. Josling GA, Russell TJ, Venezia J, Orchard L, van Biljon R, Painter HJ, et al. Dissecting the role of PfAP2-G in malaria gametocytogenesis. Nat Commun 2020 111. 2020;11: 1–13. doi:10.1038/s41467-020-15026-0

45. Painter HJ, Campbell TL, Llinás M. The Apicomplexan AP2 family: Integral factors regulating Plasmodium development. Molecular and Biochemical Parasitology. NIH Public Access; 2011. pp. 1–7. doi:10.1016/j.molbiopara.2010.11.014

46. Gissot M, Briquet S, Refour P, Boschet C, Vaquero C. PfMyb1, a Plasmodium falciparum transcription factor, is required for intra-erythrocytic growth and controls key genes for cell cycle regulation. J Mol Biol. 2005;346: 29–42. doi:10.1016/j.jmb.2004.11.045

47. Campelo Morillo RA, Tong X, Xie W, Abel S, Orchard LM, Daher W, et al. The transcriptional regulator HDP1 controls expansion of the inner membrane complex during early sexual differentiation of malaria parasites. Nat Microbiol. 2022;7: 289–299. doi:10.1038/s41564-021-01045-0

48. Tomas AM, Margos G, Dimopoulos G, Van Lin LHM, De Koning-Ward TF, Sinha R, et al. P25 and P28 proteins of the malaria ookinete surface have multiple and partially redundant functions. EMBO J. 2001;20: 3975–3983. doi:10.1093/emboj/20.15.3975

49. Dessens JT, Beetsma AL, Dimopoulos G, Wengelnik K, Crisanti A, Kafatos FC, et al. CTRP is essential for mosquito infection by malaria ookinetes. EMBO J. 1999;18: 6221–6227. doi:10.1093/emboj/18.22.6221

50. Yuda M, Sakaida H, Chinzei Y. Targeted disruption of the Plasmodium berghei CTRP gene reveals its essential role in malaria infection of the vector mosquito. J Exp Med. 1999;190: 1711–1715. doi:10.1084/jem.190.11.1711

51. Yuda M, Yano K, Tsuboi T, Torii M, Chinzei Y. von Willebrand factor A domain-related protein, a novel microneme protein of the malaria ookinete highly conserved throughout Plasmodium parasites. Mol Biochem Parasitol. 2001;116: 65–72. doi:10.1016/S0166-6851(01)00304-8

52. Langer RC, Vinetz JM. Plasmodium ookinete-secreted chitinase and parasite penetration of the mosquito peritrophic matrix. Trends in Parasitology. Elsevier Current Trends; 2001. pp. 269–272. doi:10.1016/S1471-4922(01)01918-3

53. Gerton JL, DeRisi JL. Mnd1p: An evolutionarily conserved protein required for meiotic recombination. Proc Natl Acad Sci U S A. 2002;99: 6895–6900. doi:10.1073/pnas.102167899

54. Dessens JT, Sidén-Kiamos I, Mendoza J, Mahairaki V, Khater E, Vlachou D, et al. Soap, a novel malaria ookinete protein involved in mosquito midgut invasion and oocyst development. Mol Microbiol. 2003;49: 319–329. doi:10.1046/j.1365-2958.2003.03566.x

55. Gaskins E, Gilk S, DeVore N, Mann T, Ward G, Beckers C. Identification of the membrane receptor of a class XIV myosin in Toxoplasma gondii. J Cell Biol. 2004;165: 383–393. doi:10.1083/jcb.200311137

56. Kadota K, Ishino T, Matsuyama T, Chinzei Y, Yuda M. Essential role of membrane-attack protein in malarial transmission to mosquito host. Proc Natl Acad Sci U S A. 2004;101: 16310–16315. doi:10.1073/pnas.0406187101

57. Enomoto R, Kinebuchi T, Sato M, Yagi H, Shibata T, Kurumizaka H, et al. Positive role of the mammalian TBPIP/HOP2 protein in DMC1-mediated homologous pairing. J Biol Chem. 2004;279: 35263–35272. doi:10.1074/jbc.M402481200

58. Kariu T, Ishino T, Yano K, Chinzei Y, Yuda M. CelTOS, a novel malarial protein that mediates transmission to mosquito and vertebrate hosts. Mol Microbiol. 2006;59: 1369–1379. doi:10.1111/j.1365-2958.2005.05024.x

59. Ecker A, Pinto SB, Baker KW, Kafatos FC, Sinden RE. Plasmodium berghei: Plasmodium perforin-like protein 5 is required for mosquito midgut invasion in Anopheles stephensi. Exp Parasitol. 2007;116: 504–508. doi:10.1016/j.exppara.2007.01.015

60. Tremp AZ, Khater EI, Dessens JT. IMC1b is a putative membrane skeleton protein involved in cell shape, mechanical strength, motility, and infectivity of malaria ookinetes. J Biol Chem. 2008;283: 27604–27611. doi:10.1074/jbc.M801302200

61. Ecker A, Bushell ESC, Tewari R, Sinden RE. Reverse genetics screen identifies six proteins important for malaria development in the mosquito. Mol Microbiol. 2008;70: 209–220. doi:10.1111/j.1365-2958.2008.06407.x

62. Bullen HE, Tonkin CJ, O’Donnell RA, Tham WH, Papenfuss AT, Gould S, et al. A novel family of apicomplexan glideosome-associated proteins with an inner membrane-anchoring role. J Biol Chem. 2009;284: 25353–25363. doi:10.1074/jbc.M109.036772

63. Reininger L, Tewari R, Fennell C, Holland Z, Goldring D, Ranford-Cartwright L, et al. An essential role for the Plasmodium Nek-2 Nima-related protein kinase in the sexual development of malaria parasites. J Biol Chem. 2009;284: 20858– 20868. doi:10.1074/jbc.M109.017988

64. Beck JR, Rodriguez-Fernandez IA, de Leon JC, Huynh MH, Carruthers VB, Morrissette NS, et al. A Novel Family of Toxoplasma IMC Proteins Displays a Hierarchical Organization and Functions in Coordinating Parasite Division. PLoS Pathog. 2010;6. doi:10.1371/JOURNAL.PPAT.1001094

65. Saeed S, Carter V, Tremp AZ, Dessens JT. Plasmodium berghei crystalloids contain multiple LCCL proteins. Mol Biochem Parasitol. 2010;170: 49–53. doi:10.1016/j.molbiopara.2009.11.008

66. Dessens JT, Saeed S, Tremp AZ, Carter V. Malaria crystalloids: Specialized structures for parasite transmission? Trends Parasitol. 2011;27: 106–110. doi:10.1016/j.pt.2010.12.004

67. Poulin B, Patzewitz EM, Brady D, Silvie O, Wright MH, Ferguson DJP, et al. Unique apicomplexan IMC sub-compartment proteins are early markers for apical polarity in the malaria parasite. Biol Open. 2013;2: 1160–1170. doi:10.1242/bio.20136163

68. Saeed S, Carter V, Tremp AZ, Dessens JT. Translational repression controls temporal expression of the Plasmodium berghei LCCL protein complex. Mol Biochem Parasitol. 2013;189: 38–42. doi:10.1016/j.molbiopara.2013.04.006

69. Zheng W, Liu F, He Y, Liu Q, Humphreys GB, Tsuboi T, et al. Functional characterization of Plasmodium berghei PSOP25 during ookinete development and as a malaria transmission-blocking vaccine candidate. Parasites and Vectors. 2017;10: 1–11. doi:10.1186/S13071-016-1932-4/TABLES/2

70. Deligianni E, Silmon de Monerri NC, McMillan PJ, Bertuccini L, Superti F, Manola M, et al. Essential role of Plasmodium perforin-like protein 4 in ookinete midgut passage. PLoS One. 2018;13: e0201651. doi:10.1371/journal.pone.0201651

71. Wang X, Qian P, Cui H, Yao L, Yuan J. A protein palmitoylation cascade regulates microtubule cytoskeleton integrity in Plasmodium. EMBO J. 2020;39:e104168. doi:10.15252/embj.2019104168

72. Ukegbu C V., Giorgalli M, Tapanelli S, Rona LDP, Jaye A, Wyer C, et al. PIMMS43 is required for malaria parasite immune evasion and sporogonic development in the mosquito vector. Proc Natl Acad Sci U S A. 2020;117: 7363– 7373. doi:10.1073/pnas.1919709117

73. Tachibana M, Iriko H, Baba M, Torii M, Ishino T. PSOP1, putative secreted ookinete protein 1, is localized to the micronemes of Plasmodium yoelii and P. berghei ookinetes. Parasitol Int. 2021;84:102407. doi:10.1016/j.parint.2021.102407

